# Responses of carbapenemase-producing and non-producing carbapenem-resistant *Pseudomonas aeruginosa* strains to meropenem revealed by quantitative tandem mass spectrometry proteomics

**DOI:** 10.1101/2022.10.23.513223

**Authors:** Francisco Salvà-Serra, Daniel Jaén-Luchoro, Nachiket P. Marathe, Ingegerd Adlerberth, Edward R. B. Moore, Roger Karlsson

## Abstract

*Pseudomonas aeruginosa* is an opportunistic pathogen with increasing incidence of multidrug-resistant strains, including resistance to last-resort antibiotics, such as carbapenems. Resistances are often due to complex interplays of naturally and acquired resistance mechanisms that are enhanced by its remarkably large regulatory network.

This study describes the proteomic responses of two carbapenem-resistant *P. aeruginosa* strains of high-risk clones ST235 and ST395 to subminimal inhibitory concentrations (sub-MICs) of meropenem by identifying differentially expressed proteins and pathways. Strain CCUG 51971, carries a VIM-4 metallo-β-lactamase or ‘classical’ carbapenemase, and strain CCUG 70744 carries no known acquired carbapenem-resistance genes and exhibits ‘non-classical’ carbapenem-resistance. Each strain was cultivated with different sub-MICs of meropenem, and analyzed, using quantitative shotgun proteomics, based on tandem mass tag (TMT) isobaric labeling followed by nano-liquid chromatography tandem-mass spectrometry.

Exposure of both strains to sub-MICs meropenem resulted in hundreds of differentially expressed proteins, including β-lactamases, proteins associated with transport, peptidoglycan metabolism, cell wall organization, and regulatory proteins. Strain CCUG 51971 showed up-regulation of intrinsic β-lactamases and VIM-4 carbapenemase, while CCUG 70744 exhibited a combination of up-regulated intrinsic β-lactamases, efflux pumps, penicillin-binding proteins and down-regulation of porins. All components of the H1 type VI secretion system were up-regulated in strain CCUG 51971. Enrichment analyses revealed multiple metabolic pathways affected in both strains.

Sub-MICs of meropenem cause marked changes in the proteomes of carbapenem-resistant strains of *P. aeruginosa* exhibiting different resistance mechanisms, involving a wide range of proteins, many uncharacterized, which might play a role in the susceptibility of *P. aeruginosa* to meropenem.

## Introduction

*Pseudomonas aeruginosa* is an adaptable and widely-distributed Gram-negative bacterium, which is found largely in environments associated with human activity (1, 2) and that can cause a wide range of opportunistic infections (3). *P. aeruginosa* is one of the leading causes of severe nosocomial infections, including ventilator-associated pneumonia, the most common infection among intensive care unit patients (4). It is a major cause of chronic respiratory infections in patients with cystic fibrosis or other underlying conditions such as bronchiectasis or chronic obstructive pulmonary disease (5–7).

*P. aeruginosa* is one of the most problematic drug-resistant pathogens and treatment options are often challenged by the increasing emergence of multidrug- and extensively drug-resistant strains (8, 9), which are associated with increased morbidity, as well as mortality (10–12). Indeed, *P. aeruginosa* is among the six leading pathogens causing deaths associated with antimicrobial resistance (13). Carbapenem antibiotics, such as meropenem, still remain active and effective against many multidrug-resistant isolates (9). Carbapenems are β-lactam antibiotics and thus they inhibit cell wall biosynthesis by entering the periplasmatic space and binding penicillin-binding proteins (PBPs), which are involved in the synthesis of peptidoglycan (14, 15). Compared with other β-lactams such as penicillins and cephalosporins, carbapenems have broader activity spectrum, are less susceptible to hydrolysis by β-lactamases and, in some cases, act as β-lactamase inhibitors; therefore, they are frequently used as “last-resort antibiotics” (16, 17). However, the frequency of infections caused by *P. aeruginosa* strains resistant to carbapenems is increasing worldwide (8); the World Health Organization (WHO) included carbapenem-resistant *P. aeruginosa* in the Priority 1 (critical) group of the list of priority pathogens for which research and development of new antibiotics is urgently needed (18).

In *P. aeruginosa*, carbapenem-resistance is often driven by an interplay of well-known intrinsic, acquired and adaptive resistance mechanisms, that are typically classified into three broad categories: drug transport, drug inactivation and target modification (19). These include, for instance, a naturally low outer membrane permeability, mutations leading to truncated outer membrane porins or over-expression of efflux pumps, presence of naturally-occurring and horizontally-acquired β-lactamases, and regulation or alteration of PBPs (20, 21).

However, the mechanisms behind carbapenem resistance are not always obvious (22, 23) and, for instance, mutations in core metabolic genes of *E. coli* recently have been shown to confer resistance to various antibiotics, and to be more common than previously thought. This strengthens the hypothesis that alternative non-canonical resistance mechanisms can play important roles in drug resistance (24). In fact, the presence/absence of dozens of genes, including genes involved in metabolism, have been shown to alter the susceptibility to β-lactam antibiotics in *P. aeruginosa* (25). Thus, multiple individual low-level resistance mechanisms that would not be significant on their own, collectively may contribute to the development of clinically-relevant levels of resistance. Unveiling the unknown resistance factors is essential for gaining a deeper understanding of carbapenem resistance in bacteria (26). Furthermore, this complex interplay is enhanced by the remarkably large regulatory network of *P. aeruginosa* and its high degree of responsive capacity to environmental stimuli (27, 28).

In this context, applying non-targeted open approaches, such as shotgun methodologies, is essential to understand the complexity of resistance mechanisms and the strains global responses, as well as to identify possible novel mechanisms and pathways involved in the response and resistance to antibiotics. Among those, quantitative proteomic techniques have the capacity to determine variations between conditions in the relative abundance of thousands of proteins and, therefore, such approaches have potential to reveal mechanisms involved in the responses to antibiotics that would not be possible to elucidate using classical methods of resistance detection (29, 30).

The aim of this study was to identify responses of carbapenem- and extensively drug-resistant clinical *P. aeruginosa* strains exposed to varying sub-lethal concentrations of meropenem. We further identified differentially expressed proteins (i.e., proteins with up- or down-regulated abundance levels; from now on, referred to as, “up-” or “down-regulated” proteins), groups of proteins and pathways potentially implicated in carbapenem resistance and the responses to sub-minimal inhibitory concentrations (sub-MICs) of meropenem of carbapenemase-producing (i.e., ‘classical’ carbapenem resistance) and carbapenemase non-producing (i.e., ‘non-classical’ carbapenem resistance) *P. aeruginosa* strains. For that, a “bottom-up” quantitative shotgun proteomic approach, using tandem mass tags (TMT) peptide labeling (31), followed by nano-liquid chromatography tandem-mass spectrometry (nano-LC-MS/MS) was applied.

## Materials and Methods

### Overview of the methodology

Two carbapenem-resistant *P. aeruginosa* strains, one with a ‘classical’ carbapenemase-dependent resistance (CCUG 51971) and one with a ‘non-classical’, undefined resistance (CCUG 70744) were cultivated with three different sub-MIC levels of meropenem and without antibiotic. Subsequently, proteins were extracted, reduced, alkylated and digested with trypsin, with the resulting peptides labeled, using TMTs (31). For each strain, two TMT 11-plex sets were used and four technical replicates per condition were made (two in each TMT set). *P. aeruginosa* CCUG 56489 (= PAO1), was cultivated without antibiotic and included as a control in each TMT set. The labeled samples of each set were combined and analyzed by nano-LC-MS/MS. Proteins were identified and their relative abundances determined by comparing the protein expression levels at sub-MICs and no antibiotic conditions (Figure 1).

**Figure 1.**
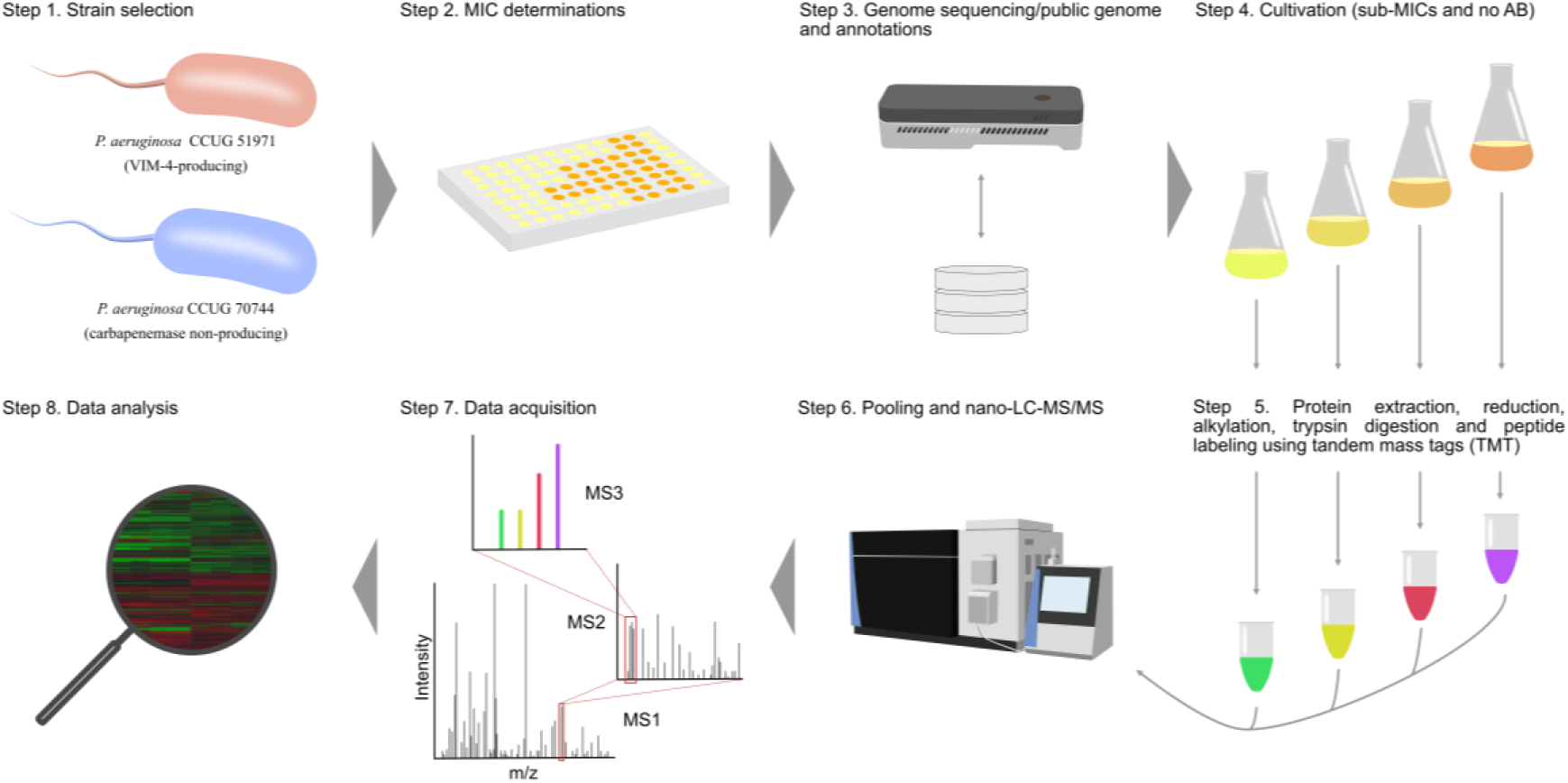
Experimental setup of the study. Each strain was cultivated with three different sub-MICs of meropenem and without antibiotic. Subsequently, proteins were extracted, reduced, alkylated and digested with trypsin. Peptides were labeled, using two TMT sets, pooled and analyzed, using nano-LC-MS/MS for protein detection and relative quantitation.

### Bacterial strains

Strain CCUG 51971 (= PA 66; sequence type 235), isolated from a human urine sample, at the Karolinska Hospital (Stockholm, Sweden) (32), carries a class-1 integron-encoded VIM-4 metallo-β-lactamase (MBL), responsible for the high carbapenem resistance levels (MIC of imipenem and meropenem >256 µg/mL; MIC of imipenem + ethylenediaminetetraacetic acid [EDTA] = 6 µg/mL) (32, 33). CCUG 51971 was the first described MBL-producing *P. aeruginosa* isolate in Scandinavia (32). Strain CCUG 70744 (sequence type 395), isolated from a human sputum sample, at the Sahlgrenska University Hospital (Gothenburg, Sweden), carries no known acquired antibiotic resistance genes (22). Additionally, strain CCUG 56489 (= PAO1) was included as a carbapenem-susceptible control. Details of the strains are presented in Table 1.

**Table 1.** General characteristics of the strains and minimal inhibitory concentrations (MICs) of the antimicrobial agents included in the “*Pseudomonas*/*Acinetobacter* standard panel” of the National Reference Laboratory for Antibiotic Resistance (Växjö, Sweden).

| Section | Features/antibiotic | *P. aeruginosa*<br>CCUG 56489 (= PAO1) | *P. aeruginosa*<br>CCUG 51971 (= PA 66) | *P. aeruginosa*<br>CCUG 70744 |
| --- | --- | --- | --- | --- |
| General information | Isolation source | Human infected wound | Human urine | Human sputum |
|  | Geographical origin | Australia | Stockholm, Sweden | Gothenburg, Sweden |
|  | Collection date | Before 1955 | 2001-11 | 2013-01-08 |
|  | NCBI RefSeq accession number | NC_002516.2 | NZ_CP043328.1 | NZ_CP023255.1 |
|  | Genome sequence length (bp) | 6,264,404 | 7,012,798 | 6,859,232 |
|  | Number of replicons | 1 | 1 | 1 |
|  | Protein sequences<sup>a</sup> | 5,572 | 6,490 | 6,399 |
|  | ANIb (%)<sup>b</sup> | 99.16 | 98.68 | 99.20 |
|  | Sequence type | 549 | 235 | 395 |
| MIC (µg/ml) | Amikacin (aminoglycoside) | 2 (S) | >32 (R) | 16 (S) |
|  | Aztreonam (monobactam) | 4 (I) | 64 (R) | >64 (R) |
|  | Cefepime (cephalosporin) | 2 (I) | >32 (R) | >32 (R) |
|  | Ceftazidime (cephalosporin) | 2 (I) | >32 (R) | >32 (R) |
|  | Ciprofloxacin (fluoroquinolone) | ≤0.12 (I) | >8 (R) | 8 (R) |
|  | Colistin (peptide antibiotic) | 1 (S) | 1 (S) | 8 (R) |
|  | Gentamicin (aminoglycoside) | 1 (IE) | >16 (IE) | 8 (IE) |
|  | Imipenem (carbapenem) | 4 (I) | >32 (R) | 16 (R) |
|  | Levofloxacin (fluoroquinolone) | 0.5 (I) | >8 (R) | 8 (R) |
|  | Meropenem (carbapenem) | 1 (S) | 512 (R) | 16 (R) |
|  | Piperacillin-tazobactam (penam) | 8 (I) | >64 (R) | >64 (R) |
|  | Tobramycin (aminoglycoside) | 0.5 (S) | >16 (R) | 2 (S) |
|  | Trimethoprim-sulfamethoxazole (diaminopyrimidine antibiotic) | 8 (R) | >16 (R) | 4 (R) |

The drug class of each antibiotic is indicated in parenthesis. EUCAST breakpoint categories are indicated in parenthesis. <sup>a</sup>Accessioned protein products annotated on the genome sequences. <sup>b</sup>ANIb with *P. aeruginosa* DSM 50071<sup>T</sup> (GenBank accession number: CP012001.1). S:
Susceptible, standard dosing regimen (green highlighting); I: Susceptible, increased exposure (yellow highlighting); R: Resistant (bold and red highlighting); IE: Insufficient evidence that the organism or group is a good target for therapy with the agent (grey highlighting).

Lyophiles of the three strains were obtained from the Culture Collection University of Gothenburg (CCUG, Gothenburg, Sweden; www.ccug.se). The strains were reconstituted on Mueller-Hinton agar (Substrate Unit, Department of Clinical Microbiology, Sahlgrenska University Hospital), at 37°C, for 24 h.

### Antibiotic susceptibility testing

Minimal inhibitory concentrations (MIC) were determined at the National Reference Laboratory for Antibiotic Resistance (Växjö, Sweden; http://www.mikrobiologi.org/referenslaboratorium), using the broth dilution method, in accordance with the EUCAST (European Committee on Antimicrobial Susceptibility Testing) recommendations and the ISO standard 20776-1 (2006). The MICs of 13 antimicrobial agents included in the “*Pseudomonas*/*Acinetobacter* standard panel” (analysis no. 25630) were determined. Additionally, strain CCUG 51971 was tested in-house for higher concentrations of meropenem, following the same recommendations. Clinical breakpoints were set, according to the EUCAST breakpoint tables v12.0 (2022) (https://www.eucast.org/clinical_breakpoints/).

### DNA extraction and whole-genome sequencing

High-molecular weight genomic DNA of *P. aeruginosa* CCUG 51971 was obtained from fresh biomass, using a previously described protocol (34). DNA integrity was verified on a TapeStation 2200 instrument (Agilent Technologies Inc., Santa Clara, CA, USA), using a Genomic DNA ScreenTape and Reagents kit (Agilent Technologies Inc.). Subsequently, isolated genomic DNA was used to prepare a standard Illumina library at Eurofins Genomics (Konstanz, Germany), with insert sizes ranging from 130 to 680 bp, following an optimized protocol and using standard Illumina adapter sequences. The genomic DNA library was sequenced, using an Illumina NovaSeq 6000 system (Illumina, Inc., San Diego, CA, USA), to generate paired-end reads of 151 bp. Genomic DNA also was used to prepare an Oxford Nanopore sequencing library, using a Rapid Barcoding Kit (SQK-RBK004; Oxford Nanopore Technologies, Ltd., Oxford, UK), following the manufacturer’s protocol. The library was sequenced on a MinION Mk1B device (Oxford Nanopore Technologies, Ltd.), using a Flow Cell FLO-MIN106 vR9.4 and the software MinKNOWN v19.05 (Oxford Nanopore Technologies, Ltd.), with the 48-h sequencing script and default parameters.

### Genome assembly and annotation

The quality of the Illumina reads was analyzed, using CLC Genomics Grid Worker v11.0.3 (Qiagen Aarhus A/S, Aarhus, Denmark). The Oxford Nanopore reads, in raw FAST5 format, were base-called, using Guppy v3.1.5 (Oxford Nanopore Technologies, Ltd.), and the quality of the sequence reads analyzed, using NanoPlot v1.26.3 (35). Subsequently, all Illumina and Nanopore reads were assembled *de novo*, using Unicycler v0.4.7 (36). The assembly was evaluated, using QUAST v5.0.2 (37), submitted to GenBank (38) and annotated, using the NCBI Prokaryotic Genome Annotation Pipeline (PGAP) v4.9 (39). The publicly available complete genome sequences of *P. aeruginosa* PAO1 (28) and *P. aeruginosa* CCUG 70744 (22) were downloaded from the NCBI Reference Sequence (RefSeq) database (40). To confirm the species identity of the strains, average nucleotide identity values based on BLAST (ANIb) (41) were calculated, using JSpeciesWS (42), between the genome sequences of the three strains and that of the type strain of *P. aeruginosa* (DSM 50071^T^; GenBank accession number: CP012001.1). Multilocus sequence typing (MLST) of the complete genome sequences was performed, using the tool MLST v2.0.4 and the MLST database v2.0.0 of the Center for Genomic Epidemiology (CGE) (43), with the *P. aeruginosa* MLST profile (44).

### Cultivation conditions for the quantitative proteomics experiments

Preinocula were prepared in 4 ml of Mueller-Hinton broth, incubated, at 37°C, overnight, with orbital shaking (250 rpm). Cells were pelleted by centrifugating at 10,000 x *g*, for 5 min, and resuspended in phosphate-buffered saline (PBS) adjusting the turbidity to MacFarland Standard 0.5 (equal to 1-2 x 10^8^ colony forming units, CFUs/ml) (45). Subsequently, 250 µl were inoculated in baffled Erlenmeyer flasks of 250 ml capacity containing Mueller-Hinton broth, to a final volume of 50 ml. The flasks were incubated, at 37°C, for 20 h, with orbital shaking (250 rpm). Strain CCUG 56489 (= PAO1) was cultivated without antibiotic. Strain CCUG 70744 was cultivated with 0, 2, 4 (¼ of MIC) and 8 µg/ml (½ of MIC) of meropenem (Sigma-Aldrich, St. Louis, MO, USA). Strain CCUG 51971 was cultivated with 0, 8, 128 (¼ of MIC) and 256 µg/ml (½ of MIC) of meropenem.

### Peptide generation and TMT labeling

Bacterial biomasses were harvested by centrifugation (10,000 x *g*, 5 min), and washed three times with 1.0 ml of PBS, by discarding the supernatant, resuspending the pellet and centrifuging at 10,000 x *g*, for 5 min. Washed bacteria were resuspended in PBS, to optical density 1.0 at a wavelength of 600 nm (OD_600_), and divided into four tubes (four technical replicates for nano-LC-MS/MS). Bacterial cells were pelleted by centrifugation at 10,000 x *g*, for 5 min, resuspended in 150 µl of PBS plus 15 µl of 20% sodium dodecyl sulfate (SDS) and transferred to small vials (200 µl) containing acid-washed glass beads (diameter: 150 – 212 μm) (Sigma-Aldrich). The cells were lysed by bead-beating, using a TissueLyser II (Qiagen, Hilden, Germany), at 25 Hz, for 5 min, and the lysates were frozen at −20°C until analysis.

Protein concentrations were determined using a Pierce™ BCA Protein Assay kit (Thermo Fisher Scientific, Waltham, MA, USA) and a Benchmark Plus Microplate Reader (Bio-Rad, Hercules, CA, USA). Bovine serum albumin solutions were used as standards. Representative references containing equal amounts from each group were prepared. Proteins were digested with trypsin, using the filter-aided sample preparation (FASP) method (46). Aliquots containing 30 μg of protein from each sample and the references were used for digestions. Briefly, samples were reduced using 100 mM dithiothreitol, at 60°C, for 30 min, transferred to 30 kDa MWCO Pall Nanosep^®^ centrifugation filters (Sigma-Aldrich), washed repeatedly using urea (8 M) and once using digestion buffer (1% sodium deoxycholate [SDC], in 50 mM triethylammonium bicarbonate, TEAB), and alkylated using 10 mM methyl methanethiosulfonate, in digestion buffer, for 30 min. Trypsin digestion was performed in digestion buffer, by adding 0.5 µg of Pierce MS grade Trypsin (Thermo Fisher Scientific) and incubating at 37°C, overnight. An additional portion of trypsin was added and incubated for two additional hours. Peptides were collected by centrifugating, at 10,000 x *g*.

Peptides were labelled, using TMT 11-plex isobaric mass tagging reagents (Thermo Fisher Scientific), following the manufacturer’s instructions. Samples were combined into two TMT-sets with two samples from each group and a reference. SDC was removed by acidifying using 10% trifluoroacetic acid. The combined sets were pre-fractionated into 20 fractions by performing basic reversed-phase liquid chromatography (bRP-LC), using a Dionex Ultimate 3000 UPLC system (Thermo Fisher Scientific) and a reversed-phase XBridge BEH C18 column (3.5 μm, 3.0 x 150 mm, Waters Corporation, Milford, MA, USA). For that, a linear gradient was created by mixing solvent A (10 mM ammonium formate buffer, pH 10.0) and solvent B (90% acetonitrile and 10% 10 mM ammonium formate, pH 10.0) at a flow rate of 400 µl/min, increasing solvent B from 3% to 40%, over 18 min, followed by an increase to 100% of solvent B, over 5 min. The fractions were concatenated into 10 fractions (1+11, 2+12, … 10+20), dried and reconstituted in 3% acetonitrile and 0.2% formic acid.

### Nano-LC-MS/MS

The fractions were analyzed using an Orbitrap Fusion™ Lumos™ Tribrid™ mass spectrometer coupled with an Easy-nLC1200 nano-liquid chromatography system (Thermo Fisher Scientific). Fractioned peptides were trapped on an Acclaim Pepmap 100 C18 trap column (100 μm x 2 cm, particle size 5 μm, Thermo Fischer Scientific) and separated in an in-house-packed analytical column (75 μm x 30 cm, particle size 3 μm, Reprosil-Pur C18, Dr. Maisch HPLC, Ammerbuch, Germany), using a linear gradient of solvent B (80% acetonitrile, 0.2% formic acid) over solvent A (0.2% formic acid) from 5% to 33%, over 77 min, followed by an increase to 100% of solvent B for 3 min, and 100% solvent B, for 10 min, at a flow rate of 300 nL/min. Precursor ion mass spectra were acquired at 120,000 resolution and MS/MS analysis was performed in a data-dependent multi-notch mode, wherein collision-induced dissociation (CID) spectra of the most intense precursor ions were recorded in the ion trap at a collision energy setting of 35 for 3 s (‘top speed’ setting). Precursors were isolated in the quadrupole using a 0.7 m/z isolation window. For fragmentation, charge states 2 to 7 were selected. Dynamic exclusion was set to 45 s and 10 ppm. For reporter ion quantitation, MS^3^ spectra were recorded at 50,000 resolution with higher-energy collisional dissociation (HCD) fragmentation at collision energy of 65, using the synchronous precursor selection.

### Proteomic Data Analysis

The data files for each TMT set were merged for identification and relative quantitation, using Proteome Discoverer v2.4 (Thermo Fisher Scientific). The search was done against the accessioned protein products annotated on the genome sequences (i.e., FAA format files) of *P. aeruginosa* strain CCUG 56489 (= PAO1) (RefSeq accession number: NC_002516.2), *P. aeruginosa* strain CCUG 51971 (RefSeq accession number: NZ_CP043328.1) or *P. aeruginosa* CCUG 70744 (RefSeq accession number: NZ_CP023255.1), using Mascot v2.5.1 (Matrix Science, London, UK) as a search engine and both databases defined in the Protein Marker node. The precursor mass tolerance was set to 5 ppm and fragment mass tolerance to 0.6 Da. Tryptic peptides were accepted with zero missed cleavage, variable modifications of methionine oxidation and fixed cysteine alkylation, TMT-label modifications of N-terminal and lysine were selected. The reference samples were used as denominator and for calculation of the ratios. Percolator was used for the validation of identified proteins. TMT reporter ions were identified in the MS^3^ HCD spectra with 3 mmu mass tolerance, and the TMT reporter intensity values for each sample were normalized on the total peptide amount. The quantified proteins were filtered at 1% false discovery rate and grouped by sharing the same sequences to minimize redundancy. Unique and razor peptides for a given protein were considered for quantification of the proteins.

### Statistical analysis

Heatmaps and principal component analysis (PCA) plots were created in R v4.0.3. The expression values were log2-transformed. The heatmaps were generated, using the *pheatmap* function. The PCAs were done using the *prcomp* function of the stats package, and plotted, using *ggplot2*.

Protein relative abundances were calculated by comparing the samples from the cultures exposed to sub-MICs of meropenem with those from the cultures exposed to no antibiotic. Only proteins with, at least, two peptide matches were considered. A Welch’s *t*-test (i.e., unequal variances *t*-test) was performed in Microsoft Excel (four technical replicates for each sub-MIC vs. four technical replicates without antibiotic). Proteins with fold changes equal to or higher than ±1.5 and *p*-values <0.05 were considered proteins with significantly different abundances (i.e., proteins with significantly up- or down-regulated abundance levels, referred to as, “up-” or “down-regulated” proteins). Clusters of significantly up- or down-regulated proteins overlapping between conditions were calculated and visualized, using BioVenn (47). For each pairwise comparison, the fold change and the *p*-value were used to generate a volcano plot (log2(fold change) vs. -log10(*p*-value)), using Microsoft Excel. Additionally, the samples from the cultures exposed to no antibiotic were compared with the samples from strain CCUG 56489 (= PAO1) to determine differences in basal protein abundances. For that inter-strain comparison, only homologous proteins sharing ≥99% of sequence identity and with 100% coverage over the longer protein were considered. The clustering and homology filtering was performed, using Cd-hit-2d (48) at CD-HIT Suite (49).

### Functional annotations and enrichment analyses

The accessioned protein products annotated on the genome sequences and available in RefSeq (40) were compared with the those of strain PAO1 using Cd-hit-2d (48) at CD-HIT Suite (49). Information from the *Pseudomonas* Genome Database (50) was added to those proteins sharing ≥95% of sequence identity and ≥95% coverage of the longest protein. The proteins were further annotated and assigned to Gene Onthology (GO) terms (51, 52), using Blast2GO (53) implemented in OmicsBox v1.3.3 (BioBam Bioinformatics S.L., Valencia, Spain). They were also annotated using InterProScan v5.54-87.0 (54) and eggNOG-Mapper v2.1.0 (55) with eggNOG v5.0.2 (56) implemented in OmicsBox v2.0.36. eggNOG mapper annotation transfers were limited to terms with experimental evidence and one-to-one orthology (i.e., prioritizing precision). Subsequently, the GO annotations were validated to remove all redundant terms based on the GO True Path Rule and taxonomically filtered, using the class *Gammaproteobacteria* (Taxonomy ID: 1236). The remaining GO terms were mapped to Enzyme Commission (EC) numbers. KEGG (Kyoto Encyclopedia of Genes and Genomes) pathways were annotated, using OmicsBox v2.0.36, via eggNOG and the assigned EC numbers (57). The accession protein products also were annotated and classified into Clusters of Orthologous Groups (COG) categories (58), using eggNOG-Mapper v2 (55), with the eggNOG v5.0 orthology resource (56) and the conditions listed above.

Antibiotic resistance genes were searched by analyzing the accessioned protein products, using the tool, Resistance Gene Identifier (RGI) v5.1.1 of the Comprehensive Antibiotic Resistance Database (CARD) v3.0.7 (59), limiting the results to “perfect” and “strict” hits only. β-lactamase types and variants were confirmed, using the Beta-Lactamase DataBase (60). Regulatory proteins were predicted, using the webserver P2RP v2.7 (61). Putative integrative and conjugative elements (ICE) were searched, using the on-line tool ICEfinder v1.0 (62). Type VI secretion systems (T6SSs) were predicted, using SecReT6 v3.0 (63). Pfam families were searched using the HMMER webserver v2.40 (64, 65). Additional functional analyses also were done, using the InterPro webserver (66). For accuracy, the annotation and information of certain proteins were checked in the *Pseudomonas* Genome Database (50), wherein orthologue searches were performed, using DIAMOND, with default parameters (i.e., query coverage and identity cutoffs: 70%, E-value: 1 x 10^-12^) (67). Frameshifted genes were confirmed by Illumina read mapping and manual examination, using CLC Genomics Grid Worker v11.0.3 (Qiagen Aarhus A/S) with default parameters.

To test if a COG category contained a significantly higher proportion of up- or down-regulated proteins and to test if any category was significantly enriched, using all quantitated proteins as a background, a Fisher’s exact test was performed, and the Benjamini-Hochberg procedure was used to control for multiple testing. Gene Set Enrichment Analyses (GSEA) of the assigned GO terms and KEGG pathways were performed, using OmicsBox v2.0.36. The enrichment statistic was “weighted (p=1)”, and 1,000 permutations were performed. The maximum and minimum gene set sizes were 500 and 15, respectively (default). Only sets with a false discovery rate (FDR) *q*-value <0.05 were deemed significantly enriched. The results were displayed in RStudio v2021.09.2, using *ggplot2* in R v4.1.2, following a previously described protocol (68).

## Results and discussion

Carbapenem-resistant *P. aeruginosa* strains are a major threat to human health (13, 18) and numerous prominent intrinsic, acquired, and adaptive mechanisms of carbapenem-resistance have been described, such as decreased outer membrane permeability, β-lactamases or efflux pumps (20, 21). However, the mechanisms behind carbapenem resistance are not always clear (22, 23), and multiple non-canonical and often low-level resistance mechanisms might contribute towards the development of clinically significant levels of resistance (24–26).

In this study, we applied a quantitative shotgun proteomic approach to study the global effect of sub-MICs of meropenem on one carbapenemase-producing and one non-carbapenemase-producing carbapenem and extensively drug-resistant *P. aeruginosa* strain, both belonging to high-risk clones: sequence type (ST) 235, the most prevalent worldwide (69), and ST395, respectively. Previous studies have analyzed the effects of β-lactams (including carbapenems) and other antibiotics on global gene expression of *P. aeruginosa* (70–72), although to the best of our knowledge, this is the first study applying a quantitative shotgun proteomics approach to determine the global proteomic responses of carbapenemase-producing and non-producing carbapenem-resistant strains of *P. aeruginosa*, belonging to high-risk clones, against exposure to sub-lethal concentrations of meropenem.

### Antibiotic susceptibility testing

Strain CCUG 51971 encodes a VIM-4 MBL, which confers high levels of resistance to imipenem and meropenem (32). *P. aeruginosa* CCUG 70744 does not harbor any known acquired carbapenemase gene but exhibited clinically significant levels of carbapenem resistance (22). Antibiotic susceptibility testing confirmed the high level of carbapenem resistance of strain CCUG 51971 (MIC of imipenem: >32 µg/ml; MIC of meropenem: 512 µg/ml) and the clinically significant resistance levels of strain CCUG 70744 (MIC of imipenem: 16 µg/ml; MIC of meropenem: 16 µg/ml). Additionally, resistance to multiple other antibiotics belonging to various classes confirmed that both strains can be classified as extensively drug-resistant, as defined by Magiorakos *et al*. (73) (Table 1).

### Whole-genome sequencing and multilocus sequence typing

Illumina sequencing of *P. aeruginosa* CCUG 51971 produced 12,783,454 paired-end reads of 151 bp (i.e., 1.93 Gb) with an average Phred quality score of 35.8. Oxford Nanopore sequencing generated 584,423 reads with an average sequence read length of 9,258 bp (i.e., 5.41 Gb), an N50 of 18,815 bp and an average Phred quality score of 9.5.

*De novo* assembly of all sequence reads produced a complete and closed circular chromosome of 7,012,798 bp with a G+C content (mol%) of 66.06%. No plasmids were detected. Annotation revealed 6,773 genes, including 83 RNA genes, 96 pseudogenes and 6,594 protein coding sequences. The ANIb between the genome sequences of strains CCUG 51971, CCUG 70744, CCUG 56489 (= PAO1) and that of *P. aeruginosa* DSM 50071^T^ ranged from 98.68 to 99.20%, confirming their species identity (Table 1). MLST analysis revealed that *P. aeruginosa* CCUG 51971 and CCUG 70744 belong to two high-risk clones: ST235 and ST395, respectively.

### Subminimal inhibitory concentrations of meropenem induce extensive changes in the proteome

To evaluate the effect of different sub-MICs of meropenem at the proteome level, the carbapenem-resistant strains *P. aeruginosa* CCUG 51971 and CCUG 70744 were cultivated with three different concentrations of meropenem (CCUG 51971 with 8, 128 and 256 µg/ml; CCUG 70744 with 2, 4 and 8 µg/ml) and without antibiotic. After sample processing and protein digestion, peptides were labeled with TMTs and analyzed using nano-LC-MS/MS.

The analysis detected 3,603 proteins (55.5% of the accessioned protein sequences) in *P. aeruginosa* CCUG 51971; of those, 90.8% (3,271 proteins) were quantitated. In *P. aeruginosa* CCUG 70744, 3,512 proteins were detected (54.9% of the accessioned protein sequences); of those, 92.9% (3,263 proteins) were quantitated (Supplementary table 1). All detected proteins (i.e., proteins with >1 peptide matches) are listed in Supplementary tables 2 and 3, together with their relative abundances (if quantitated) and functional annotations.

Heatmaps and PCA plots (Supplementary figures 1 and 2) revealed that cultures of both strains exposed to the lowest sub-MIC and to no antibiotic, demonstrated similar protein expression levels. In both cases, the samples from the cultures exposed to the highest sub-MIC demonstrated the highest difference in protein expression levels.

The relative abundances of the identified proteins were analyzed, by comparing the samples from the cultures exposed to sub-MICs of meropenem, with those from the cultures exposed to no antibiotic. Samples from cultures exposed to higher concentrations of meropenem, had higher numbers of differentially expressed proteins and, on average, higher fold changes (Figure 2 and Supplementary figure 3). In both cases, hundreds of proteins were up-regulated and, concomitantly, hundreds down-regulated when the strains were cultivated with the highest sub-MICs (i.e., CCUG 51971 with 256 µg/ml and CCUG 70744 with 8 µg/ml, which in both cases is ½ of MIC). Hundreds of proteins were also differentially expressed in each strain when cultivated with 128 µg/ml and 4 µg/ml, respectively (i.e., ¼ of MIC), while fewer were observed for the lowest concentrations, especially in the case of *P. aeruginosa* CCUG 70744, where only six proteins were differentially expressed at 2 µg/ml. However, this does not come as a surprise, since *P. aeruginosa* has an exceptionally large regulatory network. For instance, *P. aeruginosa* PAO1 harbors 521 genes (9.4% of all annotated genes) encoding transcriptional regulators or two-component systems, which reflects its high versatility and adaptability (28). Additionally, β-lactam antibiotics have been previously shown to affect global gene expression in *P. aeruginosa* (70, 71). Other studies performing quantitative shotgun proteomic analyses of *P. aeruginosa*, in those cases exposed to hypoxic stress and silver nanoparticles, also found hundreds of differentially expressed proteins (74, 75).

**Figure 2.**
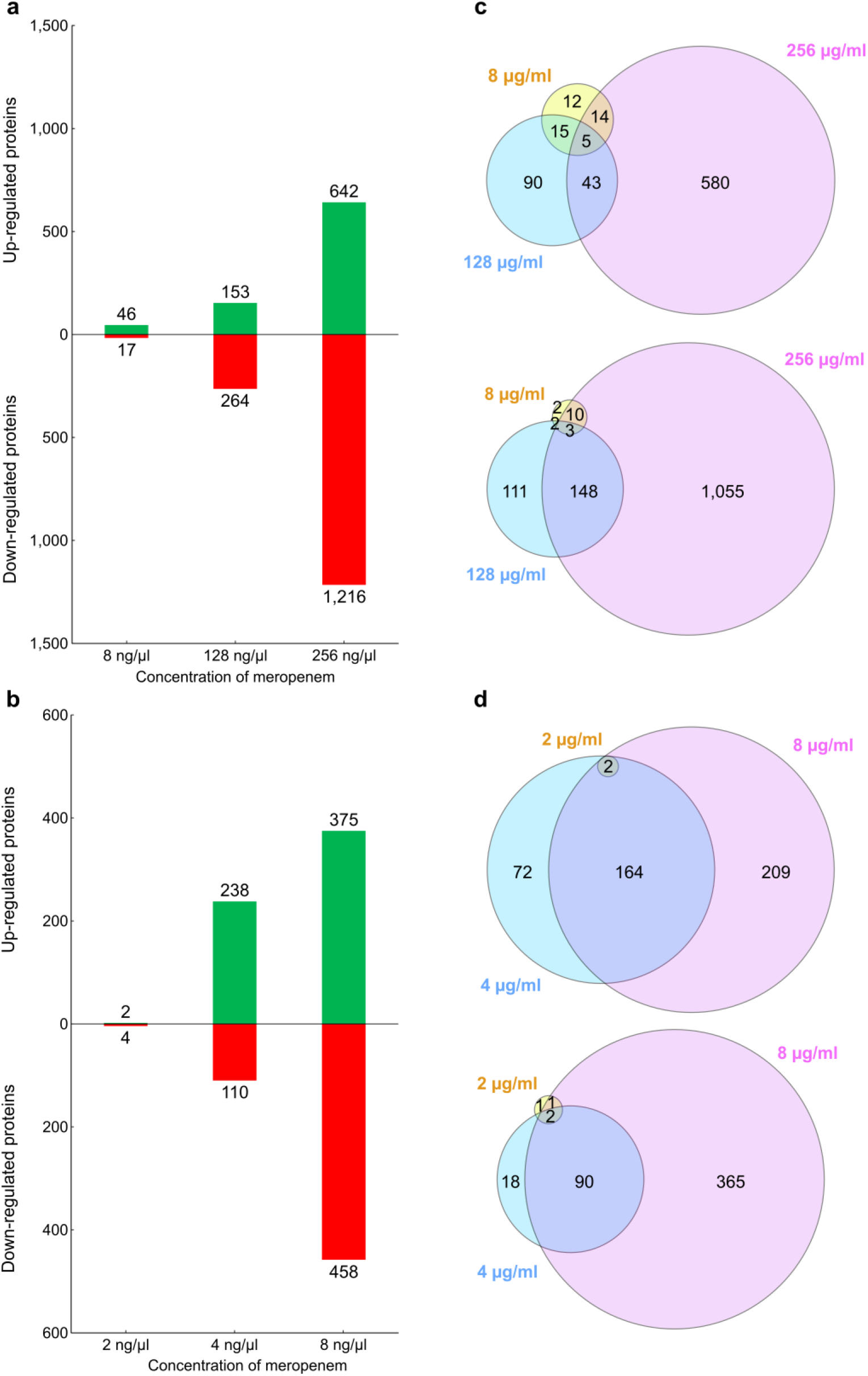
**a** Bar graph showing the number of proteins of *P. aeruginosa* CCUG 51971 up- or down-regulated in each condition. **b** Bar graph showing the number of proteins of *P. aeruginosa* CCUG 70744 up- or down-regulated in each condition. **c** Venn diagram showing the number of up- or down-regulated proteins overlapping between conditions in strain CCUG 51971. **d** Venn diagram showing the number of up- or down-regulated proteins overlapping between conditions in strain CCUG 70744.

Additionally, the samples from strains CCUG 51971 and CCUG 70744 exposed to no antibiotic were compared with the samples from strain CCUG 56489 (= PAO1) to determine differences in basal abundances. For strain CCUG 51971, a total of 2,573 proteins fulfilled the criteria for being included in the comparison (i.e., they were detected with >1 peptide, quantitated and shared >99% identity and 100% coverage with their corresponding homologue in PAO1). Of those, 335 (13%) and 423 (16%) presented higher and lower basal relative abundances, respectively (Supplementary Table 2). For strain CCUG 70744, 2,857 proteins fulfilled the inclusion criteria, of which 403 (14%) and 459 (16%) presented significantly higher and lower basal relative abundances, respectively (Supplementary Table 3). The fact that, in both strains, almost 30% of the proteins that were compared with strain PAO1 presented significantly higher or lower basal relative abundances could be expected, considering that they are different strains with a different genomic context, but establishes a base for future studies aiming to confirm them and determine the possible implications for antimicrobial resistance.

### Differentially expressed β-lactamases

*P. aeruginosa* harbors several naturally-occurring (76–78) and sometimes horizontally-acquired β-lactamases with various degrees of hydrolytic activity (79). To elucidate involvements of β-lactamases in the response to carbapenems in the two carbapenem-resistant strains included in this study, we looked at the relative abundances of these enzymes under the different sub-MICs of meropenem (Table 2).

**Table 2.**
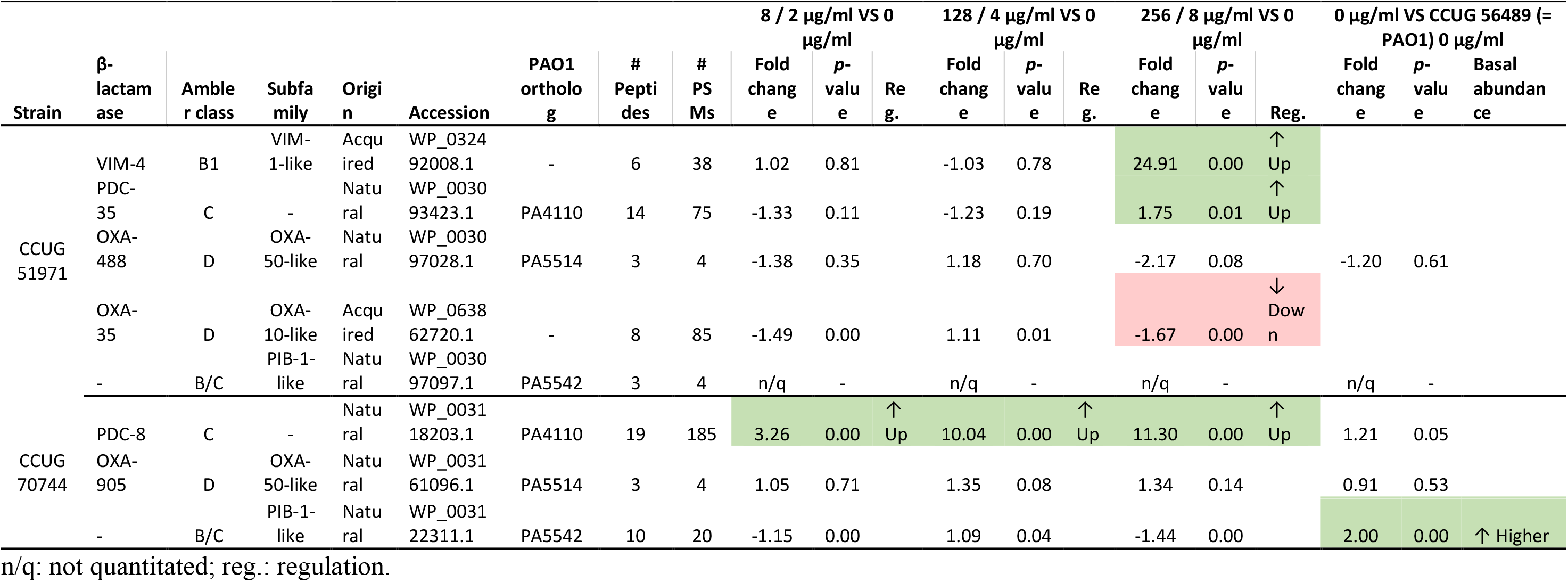
β-lactamases encoded by *P. aeruginosa* strains CCUG 51971 and CCUG 70744 and their relative abundances.

For strain CCUG 51971, a previous study showed that a class 1 integron-encoded VIM-4 MBL was responsible for its high levels of carbapenem-resistance (32). The analysis of the complete genome sequence, using ICEfinder, indicates that VIM-4 is encoded in a putative ICE of 100 kb with a type IV secretion system. The quantitative proteomic analysis showed no significant changes in the relative abundance of VIM-4 in cultures containing 8 and 128 µg/ml, suggesting that the basal expression levels of *bla*_VIM-4_ are enough for the strain to thrive in these concentrations. However, at 256 µg/ml, VIM-4 was highly up-regulated, with a fold change of 24.91, suggesting that the cells were struggling to cope with 256 µg/ml of meropenem.

Both strains encode a naturally-occurring and inducible AmpC-type Ambler class C β-lactamase (*Pseudomonas*-derived cephalosporinase, PDC). Strain CCUG 51971 encodes a PDC-35 β-lactamase, which was up-regulated at 256 µg/ml. Strain CCUG 70744 encodes a PDC-8 β-lactamase, which was up-regulated at all three sub-MICs (fold changes: 3.26, 10.04 and 11.30). Both PDC variants have an alanine residue at position 105, which has been previously associated with carbapenemase activity and reduced susceptibility to imipenem (80). Thus, the up-regulation of these PDCs with possible extended-spectrum activity might contribute to the carbapenem-resistance of these strains.

Each carbapenem-resistant strain also encodes a naturally-occurring OXA-50-like class D β-lactamase (OXA-488 and OXA-905, respectively). In both cases, the enzyme was detected and quantitated but remained stable in all conditions, which is in accordance with previous studies that failed to induce *bla*_OXA-50_, using β-lactams (77, 81). However, like OXA-50, they may have weak carbapenem hydrolytic activity (77, 81). In addition, strain CCUG 51971 also harbors a class 1 integron-located OXA-10-like restricted-spectrum oxacillinase (OXA-35), which appeared slightly down-regulated at the highest sub-MIC (-1.67-fold) and which previously has been shown to be inhibited by imipenem (82).

Finally, both carbapenem-resistant strains harbor a naturally-occurring PIB-1-like Zn^2+^-dependent imipenemase. In strain CCUG 51971, the protein was detected, but could not be quantitated, probably because it was produced at levels too low to measure. In strain CCUG 70744, the same protein was detected and quantitated, but not differentially expressed in any condition. However, the relative abundance of PIB-1 in strain CCUG 70744 was double than the one observed in strain PAO1, which has been previously suggested to contribute to carbapenem resistance (78).

### Differentially expressed porins, efflux pumps and other transport-related proteins

In Gram-negative bacteria, β-lactam antibiotics need to cross the outer membrane via porins (i.e., water-filled protein channels) in order to reach the target PBPs, which are located on the surface of the cytoplasmatic membrane (83). By default, the outer membrane permeability of *P. aeruginosa* is 12 – 100-fold lower than that of *E. coli* (84, 85), partly due to the lack of non-specific general diffusion porins and to narrower specific porins (86, 87). Since porins can be involved in the import of antibiotics, we looked for porins that were differentially expressed when the carbapenem-resistant strains were exposed to sub-MICs of meropenem.

Firstly, we looked at the outer membrane porin OprD, which mainly serves for the transport of nutrients, such as basic amino acids, but also aids the intake of carbapenems and is considered a major resistance determinant (88). In strain CCUG 51971, the genome sequence revealed an insertion causing a frameshift (1206insC; locus tag: FZE14_RS23605) and an altered 57 amino acids longer version of the protein. The quantitative proteomic analysis revealed only residual abundance ratios in all four conditions, suggesting that the protein is either not produced or produced at very low levels. In strain CCUG 70744, OprD was produced, but down-regulated at the two highest sub-MICs, showing the largest fold change at the highest sub-MIC (Table 3). Additionally, inactivation of OprD also has been associated with enhanced fitness and virulence (89), indicating that this could also led to increased pathogenicity.

**Table 3.**
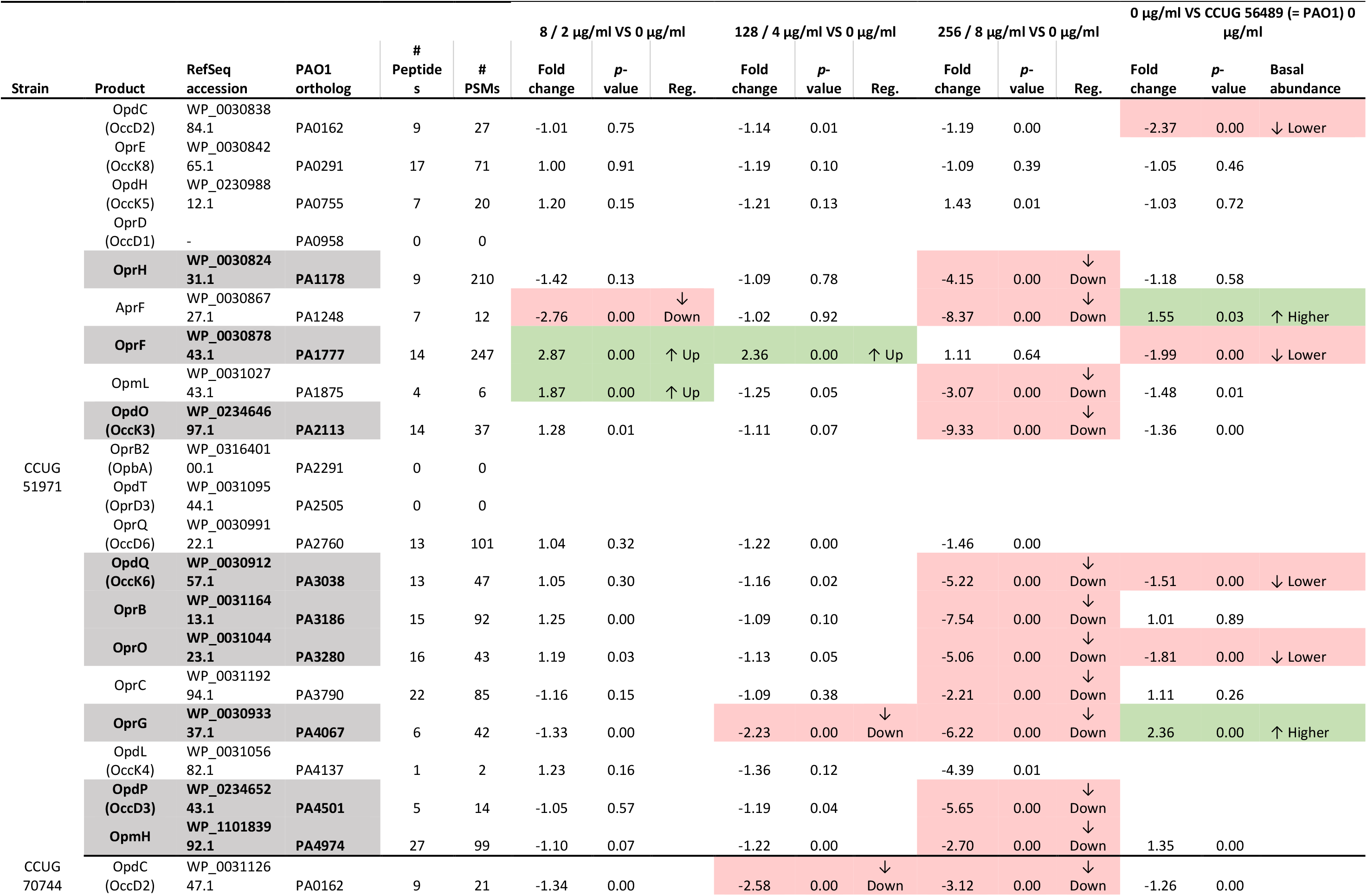
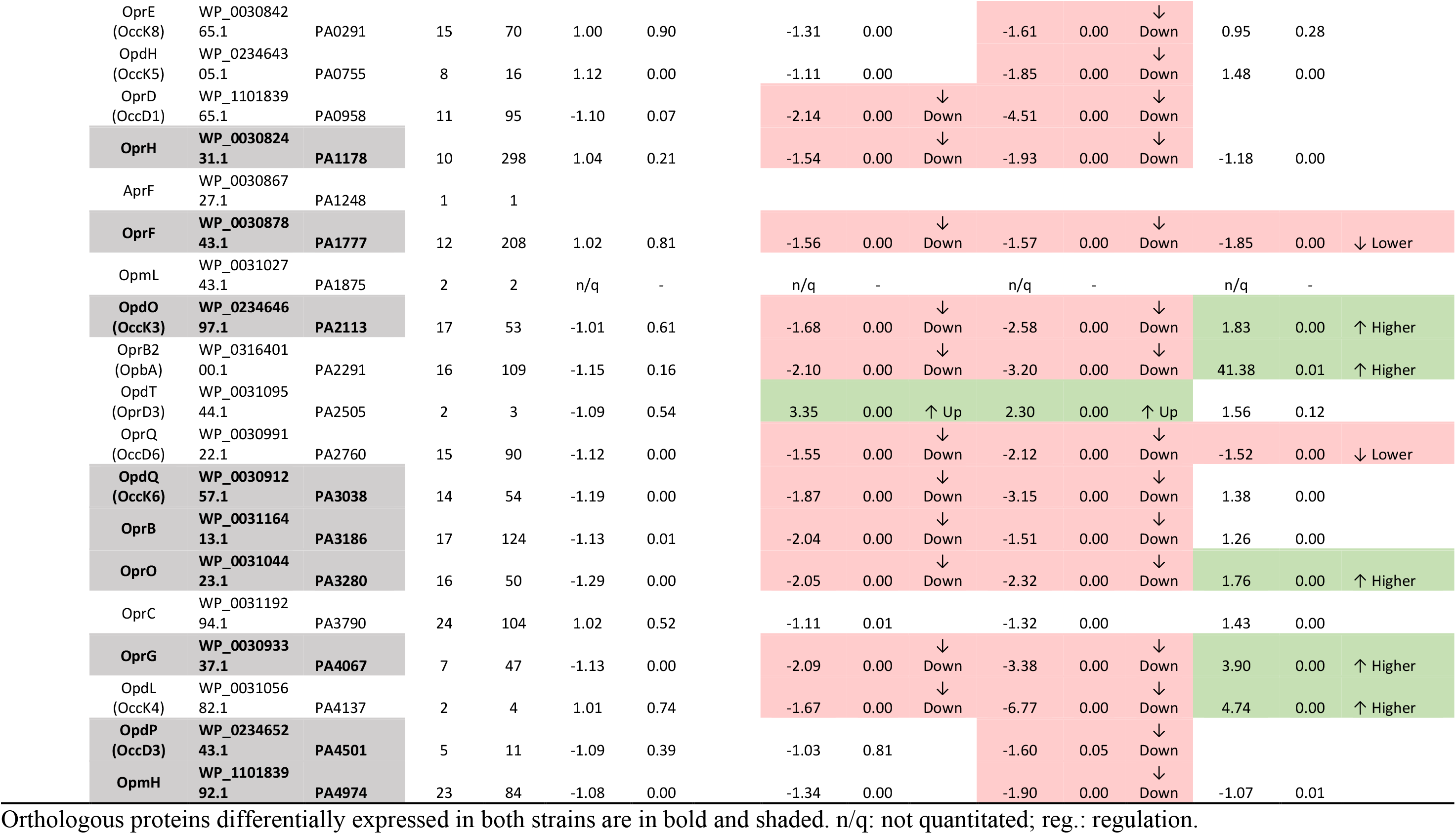
Differentially expressed porins of *P. aeruginosa* CCUG 51971 and CCUG 70744.

The genome sequence of strain CCUG 51971 also revealed that the gene encoding OpdD, another porin which can facilitate uptake of meropenem (90), was frameshifted by a deletion (213del; locus tag: FZE14_RS22945) that generates a premature stop codon. In strain CCUG 70744, OpdD seemed functional in the genome sequence but was not detected by the proteomic analysis, which could be due to very low abundances.

In both carbapenem-resistant strains, the majority of porins that were deemed differentially expressed when exposed to sub-MICs of meropenem, were down-regulated. In strain CCUG 51971, 11 porins were down-regulated at the highest sub-MIC (Table 3). Of those, OprG was also down-regulated at the middle sub-MIC and AprF at the lowest. Strikingly, OprF appeared up-regulated at 8 and 128 µg/ml and OpmL up-regulated at 8 µg/ml, but none of them at 256 µg/ml. In strain CCUG 70744, 16 porins were down-regulated at the highest sub-MIC, four of them only in that condition and 11 both at the highest and the middle sub-MICs. Interestingly, OpdT was up-regulated at 4 and 8 µg/ml. Of all these porins, eight were down-regulated in both strains (OprH, OpdP, OpdO, OpdQ, OprB, OprO, OprG and OpmH), suggesting that they might be conserved responses to meropenem in *P. aeruginosa*. Of these, OpdP has been demonstrated to be involved in meropenem uptake (90).

Additionally, *P. aeruginosa* harbors a large arsenal of efflux pumps, some of which have been associated with natural and acquired antibiotic resistance caused by extrusion of antibiotics to the outside of the cells (91). Hence, we searched for differentially expressed components of the characterized drug efflux pumps listed by Li and Plésiat (2016) (91) (Table 4).

**Table 4.**
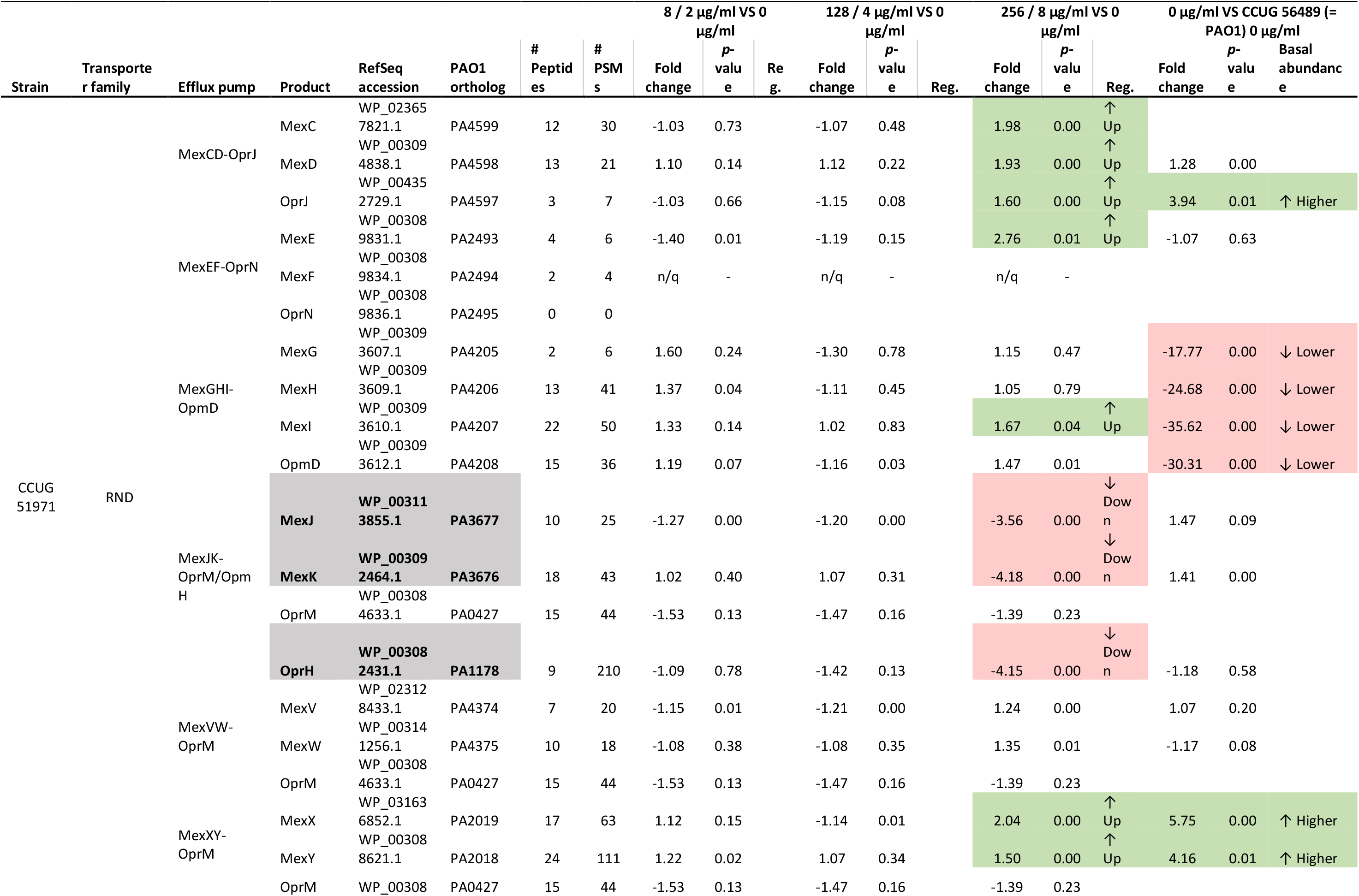
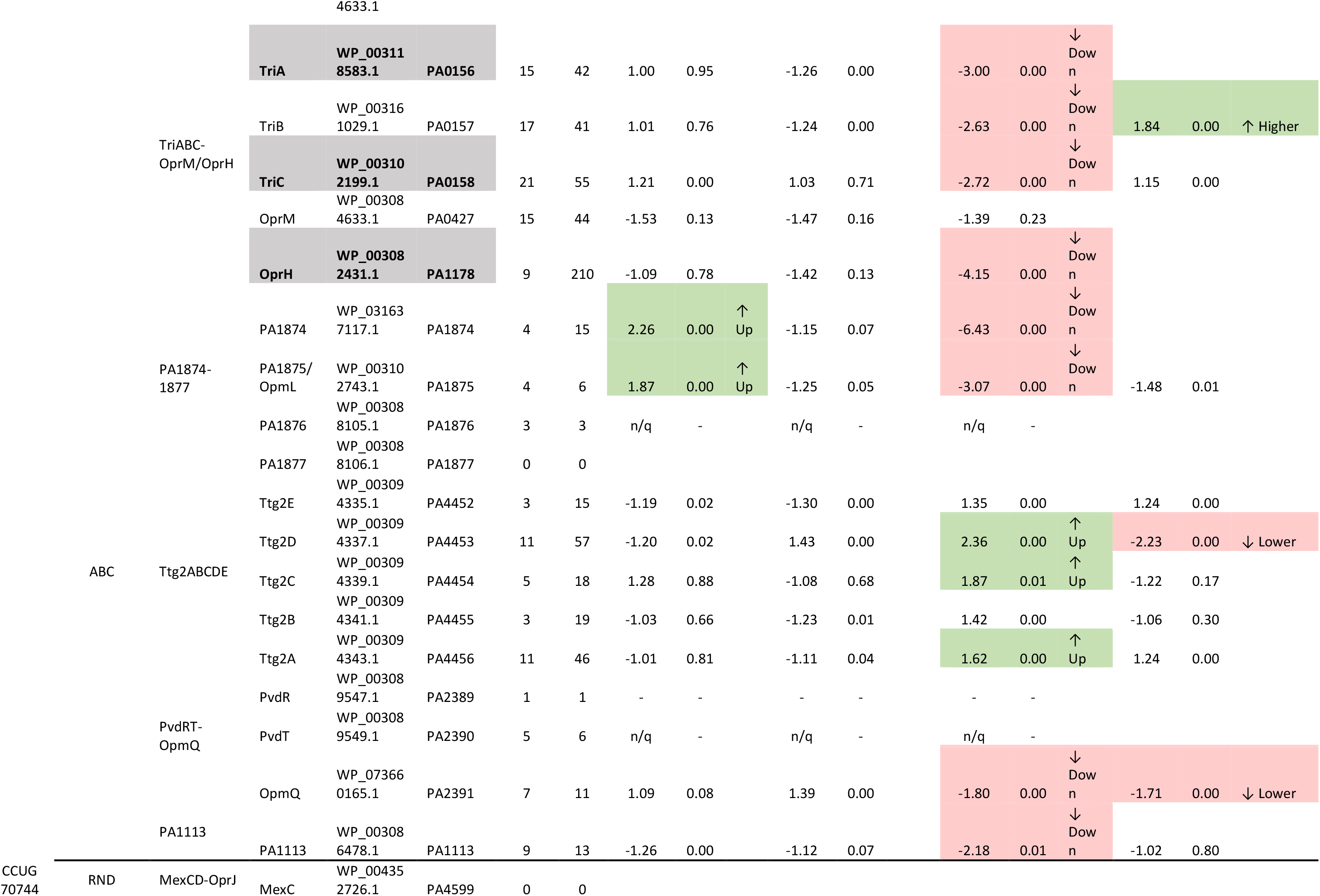
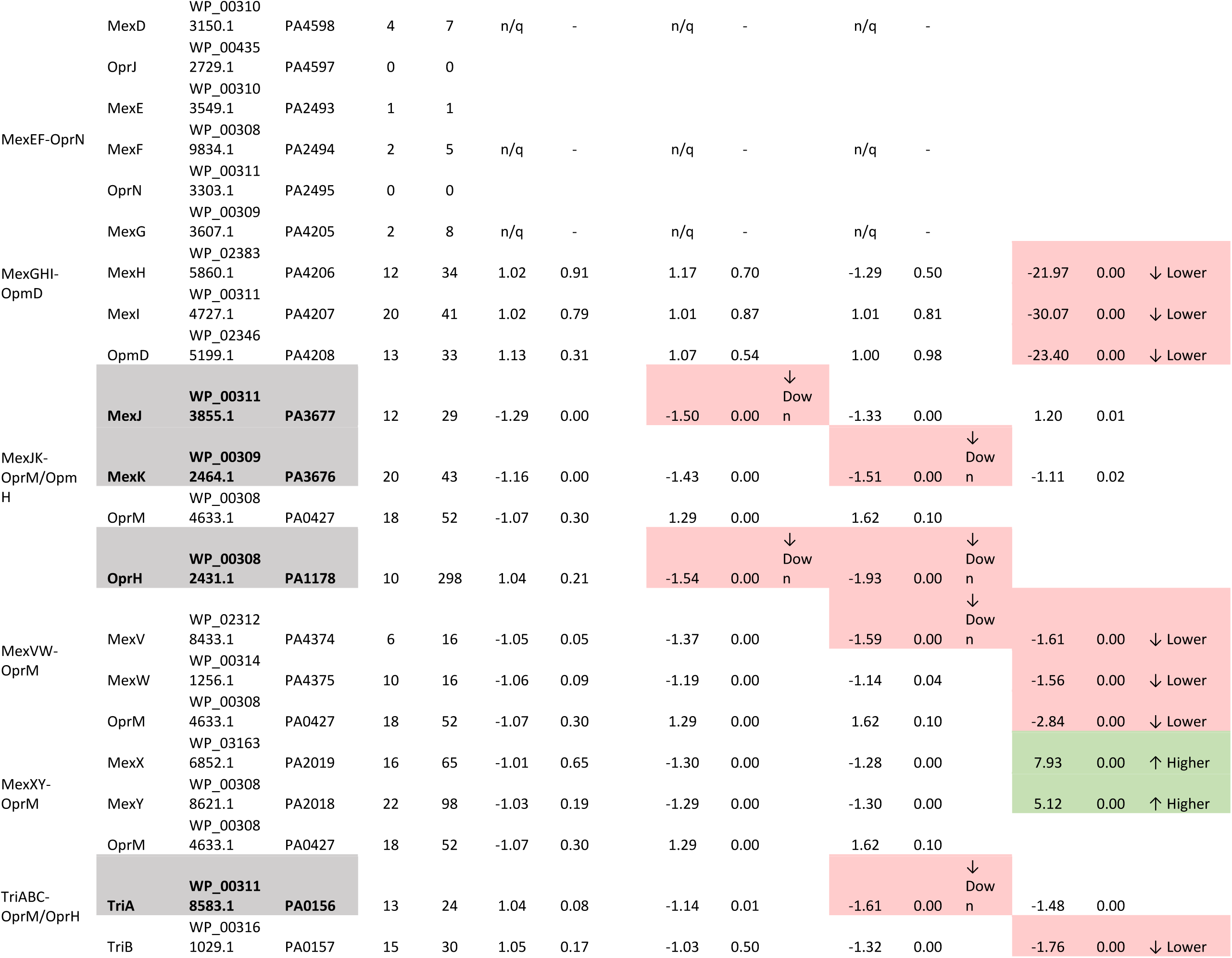
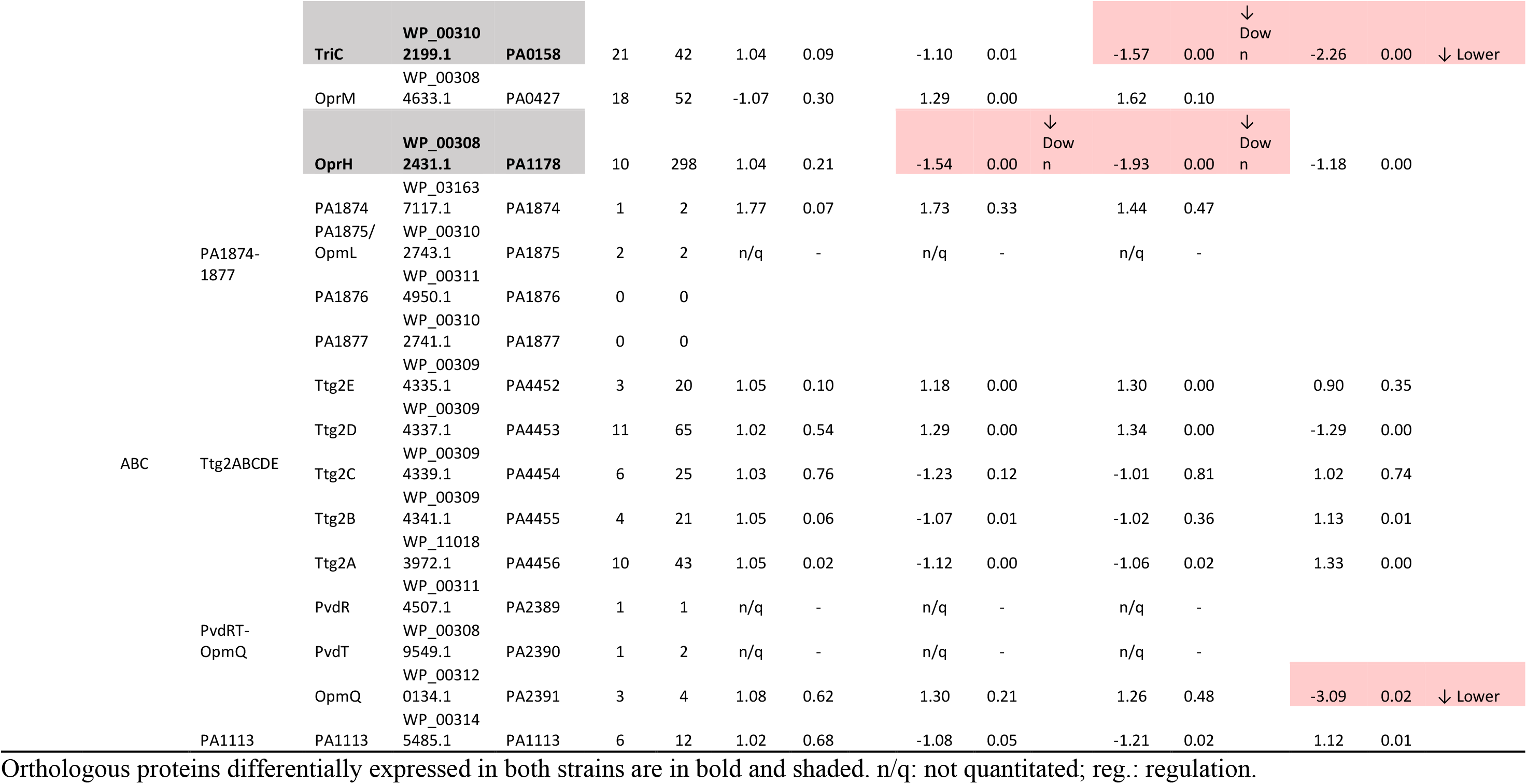
Characterized drug efflux pumps of *P. aeruginosa* CCUG 51971 and CCUG 70744 with differentially expressed proteins.

In strain CCUG 51971, several efflux pumps of the resistance-nodulation-cell division (RND) family were up-regulated at 256 µg/ml. Among those, were included the three components of the MexCD-OprJ efflux pump, which can extrude meropenem (92). The protein MexE of the MexEF-OprN efflux pump was also up-regulated, although the MexF and OprN components were not quantitated nor detected, respectively. The MeXY components of the MexXY-OprM pump, which is also able to pump out meropenem (92), also were up-regulated. Moreover, compared to strain PAO1, the basal relative abundances of the MexXY components were already 4 – 6-fold higher. Finally, the MexI component of the MexGHI-OpmD efflux pump was also up-regulated, although not the other components. Additionally, the proteins Ttg2A, Ttg2C and Ttg2D of the Ttg2 ATP-binding cassette (ABC) transporter, also were up-regulated. This transporter is associated with membrane-phospholipid homeostasis, outer membrane permeability and has been previously associated to low level resistance to various relatively hydrophobic antibiotics (93).

In contrast, all components of the MexJK-OpmH and TriABC-OprH RND efflux pumps were down-regulated at 256 µg/ml, as well as the outer membrane protein of the PvdRT-OpmQ ABC transporter. Interestingly, the two components of the ABC transporter encoded by the *P. aeruginosa* locus tags PA1874-1877 that were quantitated, appeared down-regulated at 256 µg/ml but up-regulated at 8 µg/ml.

In strain CCUG 70744, fewer efflux pumps had differentially expressed protein subunits. Like in strain CCG 51971, the RND efflux pumps MexJK-OprH and TriABC-OprH had down-regulated components (MexJ at 4 µg/ml; MexK, TriA and TriC at 8 µg/ml, and OprH at 4 and 8 µg/ml). Additionally, the protein MexV of the MexVW-OprM efflux pump was slightly down-regulated. The MexAB-OprM efflux pump has been shown to extrude meropenem (92); however, this system was not differentially expressed in any of the strains and did not present higher protein abundances compared to strain PAO1, although the three proteins of the system were detected in both cases with more than ten peptides. The MexXY-OprM efflux pump was not differentially expressed with meropenem exposure. However, when compared with strain PAO1, the MeXY components had a significantly higher relative abundance (MexX: 7.93-fold; MexY: 5.12-fold), as observed also in strain CCUG 51971. This confirms the prediction made by Johnning *et al*., who reported an insertion impairing the function of the repressor *mexZ* in the genome of strain CCUG 70744 (22), which has been previously linked to overexpression of MeXY (94), which may contribute to the low meropenem susceptibility of the strain (93).

Since lowering the outer membrane permeability and increasing efflux pumps expression are major mechanisms of resistance in *P. aeruginosa*, we searched for additional transmembrane transport-related proteins that were differentially expressed. Using several selected GO terms, 55 and 22 proteins were found to be potentially associated with transmembrane transport and differentially expressed in strains CCUG 51971 and CCUG 70744, respectively (Table 5). Most of those proteins were down-regulated (40 of 55 and 20 of 22) and 14 were differentially expressed in both strains, suggesting that they might be involved in conserved responses to meropenem.

**Table 5.**
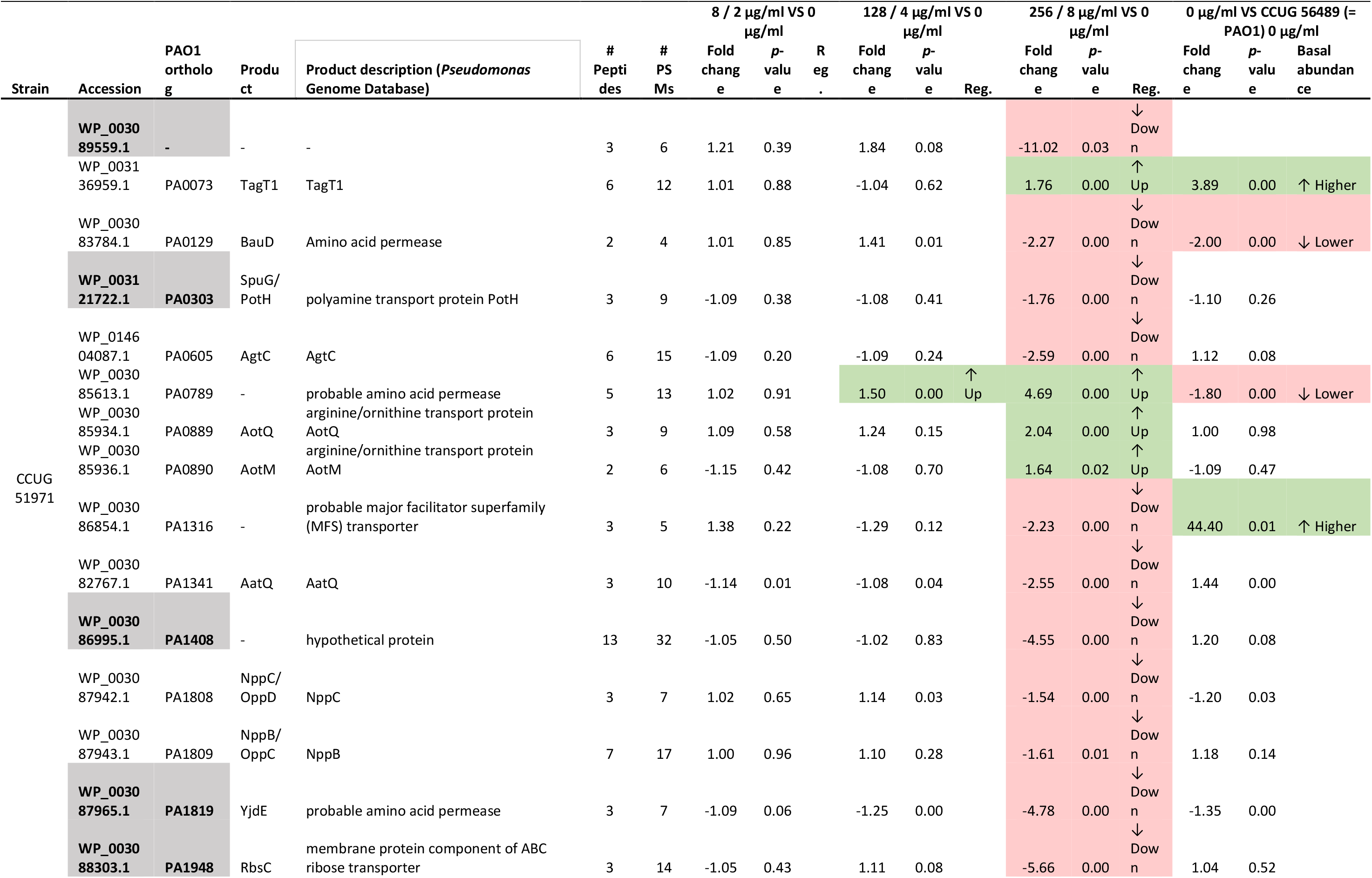
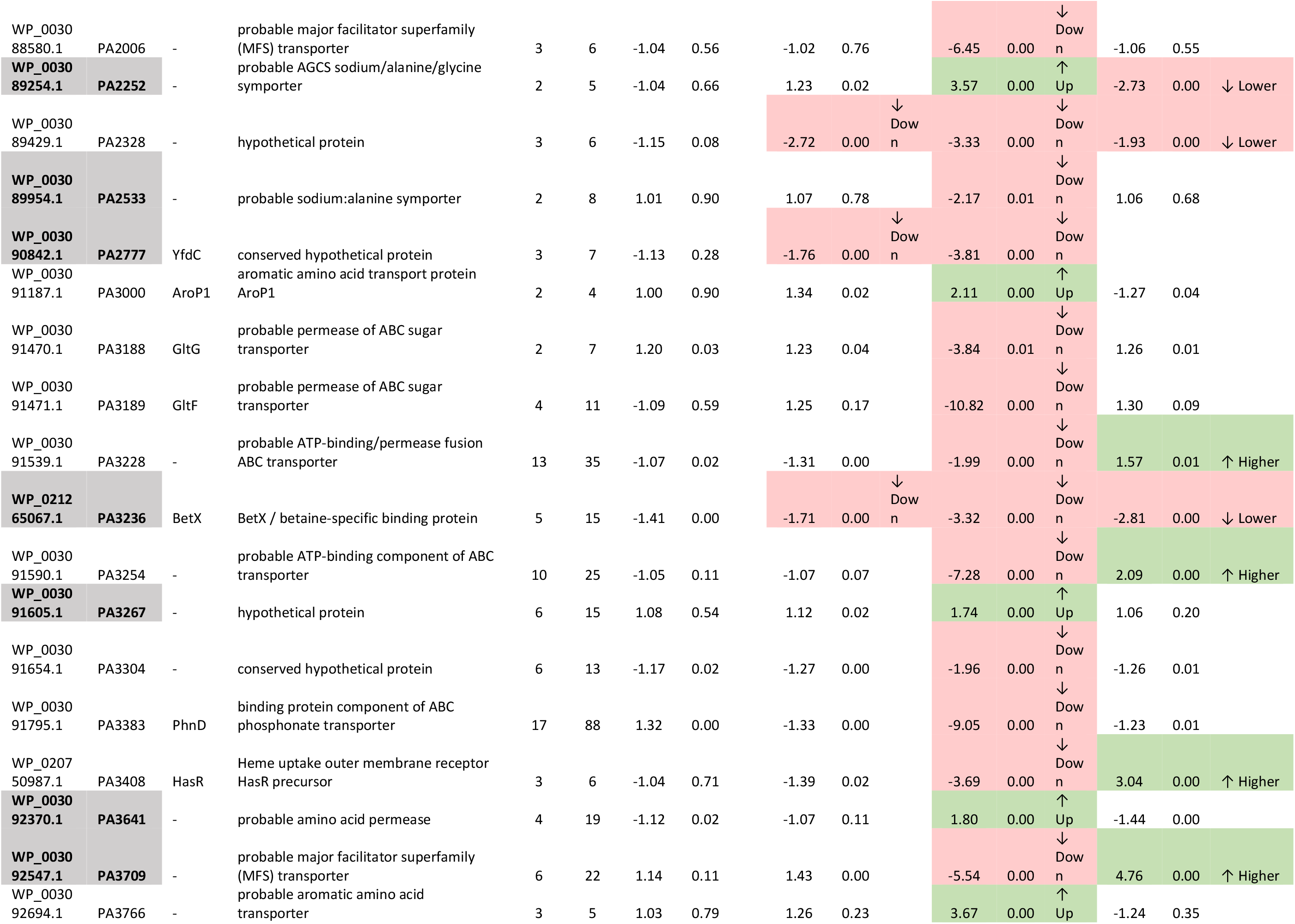
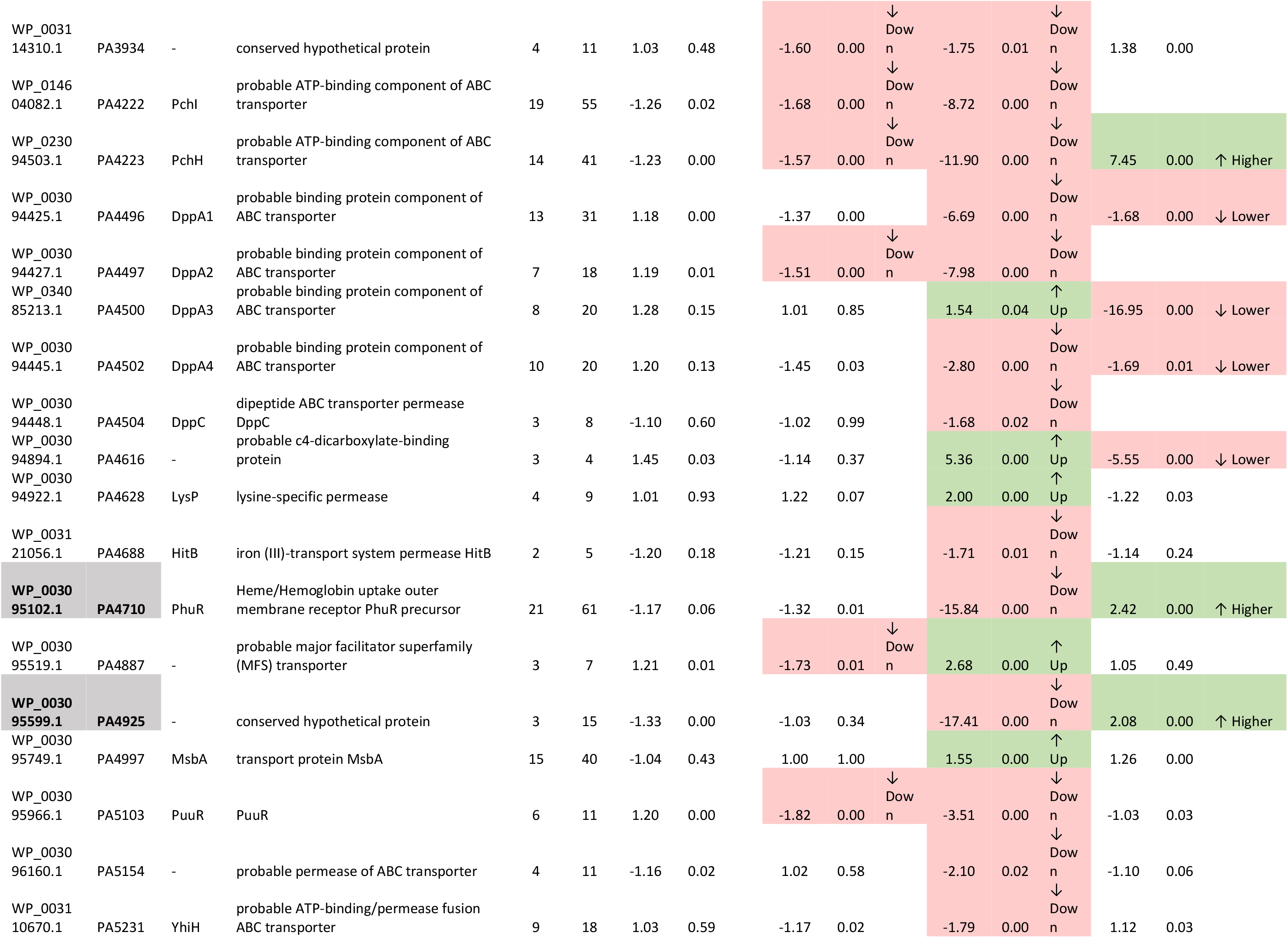
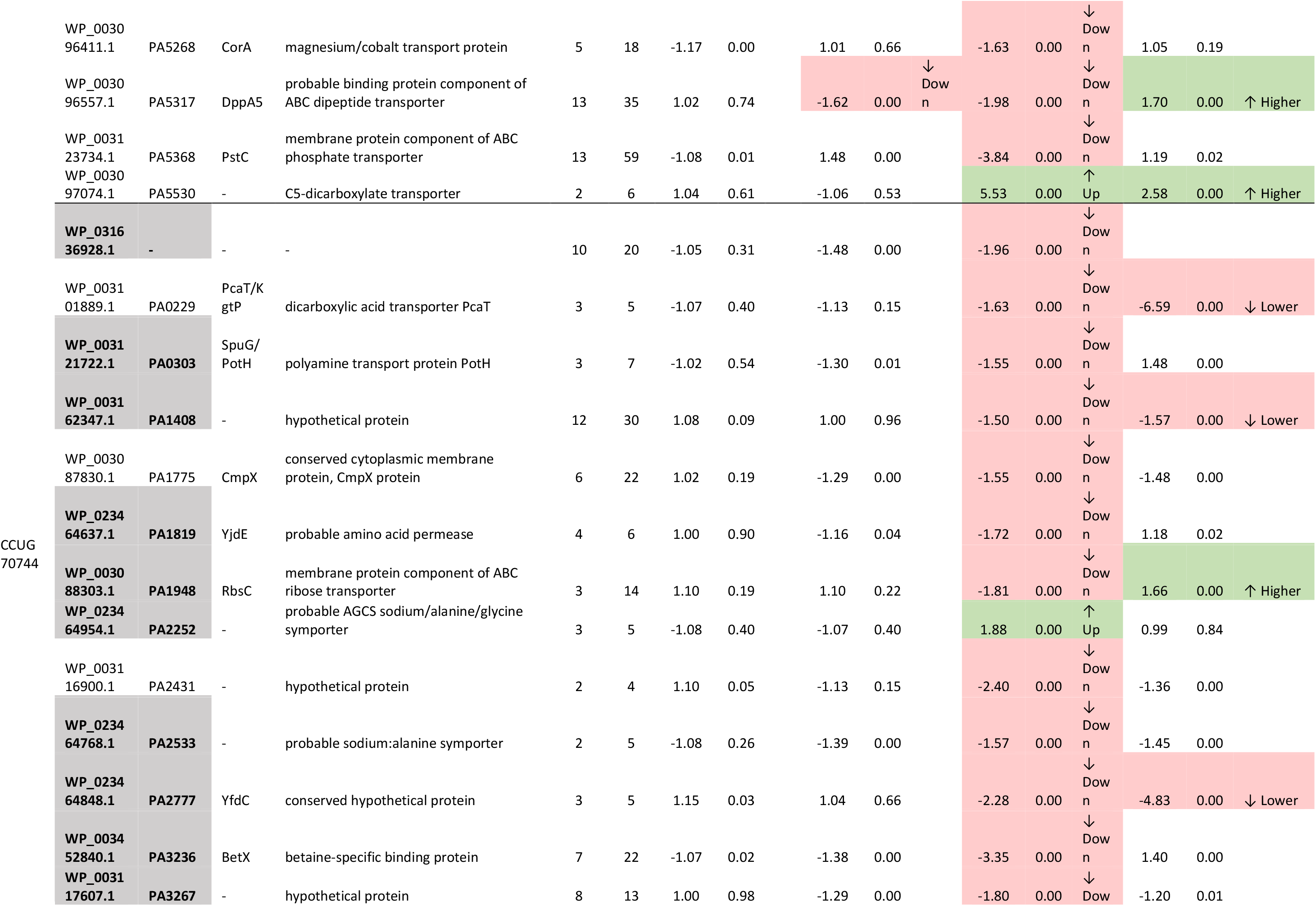
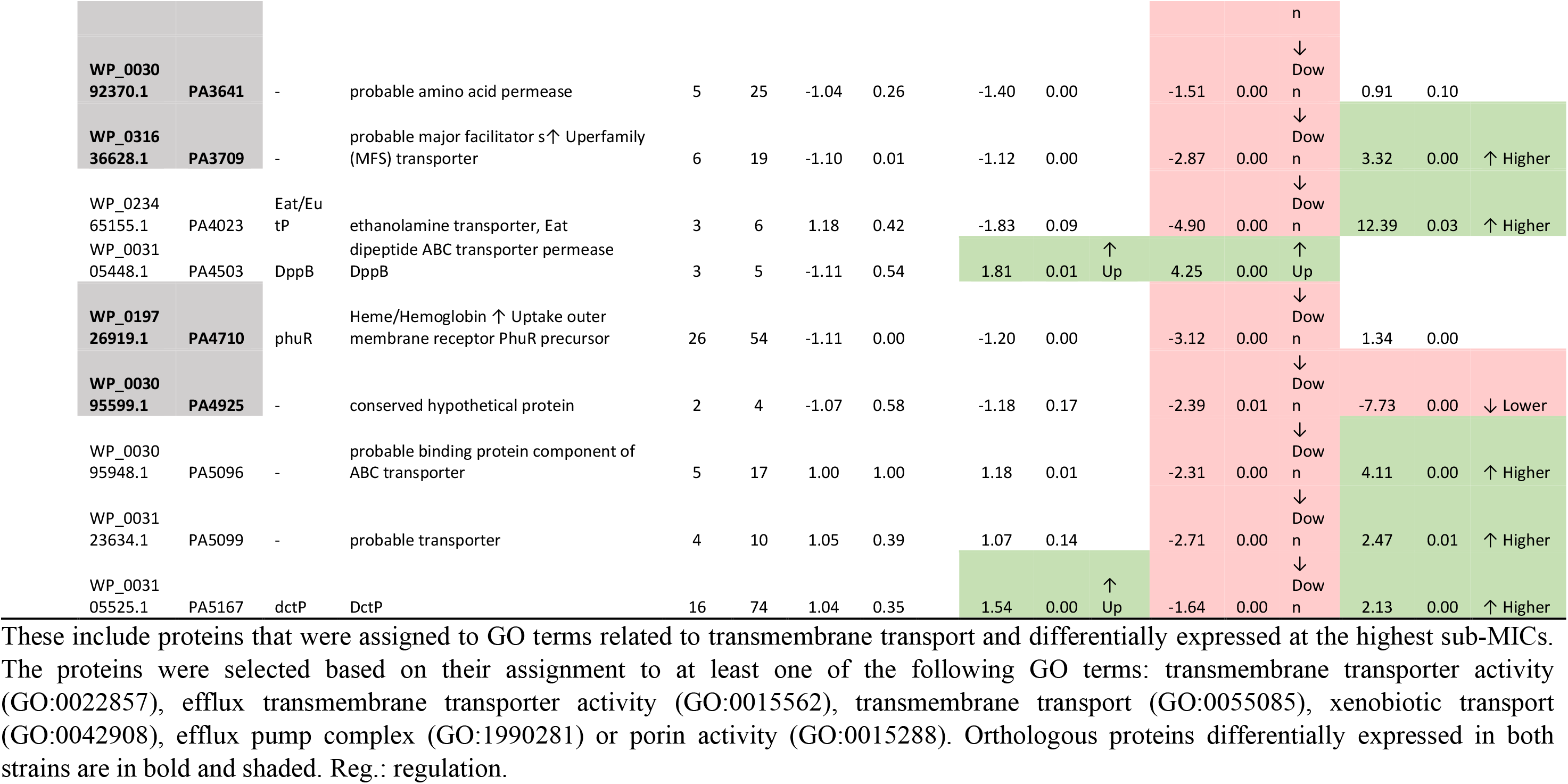
Additional differentially expressed proteins related with transmembrane transport.

### Differentially expressed penicillin-binding-proteins

PBPs, which are essential for the synthesis and maintenance of the peptidoglycan cell wall, are the targets of β-lactam antibiotics (83). Therefore, we looked at these proteins to determine which ones were differentially expressed in response to sub-MICs of meropenem in these two carbapenem-resistant strains (Table 6). In *P. aeruginosa*, at least eight PBPs have been described (95), of which two were differentially expressed, in both strains. Among those two was PBP3, one of the primary targets of meropenem (14, 96), which previously has been shown to be indispensable for *P. aeruginosa* growth (97). PBP3 was up-regulated in both strains, although at different levels. In strain CCUG 70744, PBP3 was moderately up-regulated at the two highest sub-MICs (fold changes lower than 2). In strain CCUG 51971, PBP3 was also up-regulated, but only at the highest sub-MIC and at a higher level (8.72-fold), which might be due to the shielding effect of the VIM-4 MBL (responsible for the high carbapenem resistance levels of this strain) (32) at the two lowest sub-MICs. Additionally, in strain CCUG 70744, PBP3 had a higher basal abundance, compared to that of strain PAO1. Up-regulation of PBP3 might contribute to the low susceptibilities of these strains, as overproduction of PBP3 in *P. aeruginosa* has been shown to reduce susceptibility to multiple β-lactam antibiotics (98). In fact, mutations in PBP3 also have been associated with altered β-lactams susceptibilities, which highlights its relevance (99). A similar trend also was observed for PBP7, which in strain CCUG 70744 was moderately up-regulated at the two highest sub-MICs (1.51 and 2.15-fold) while in strain CCUG 51971 it was highly up-regulated but only at the highest sub-MIC (11.75-fold).

**Table 6.**
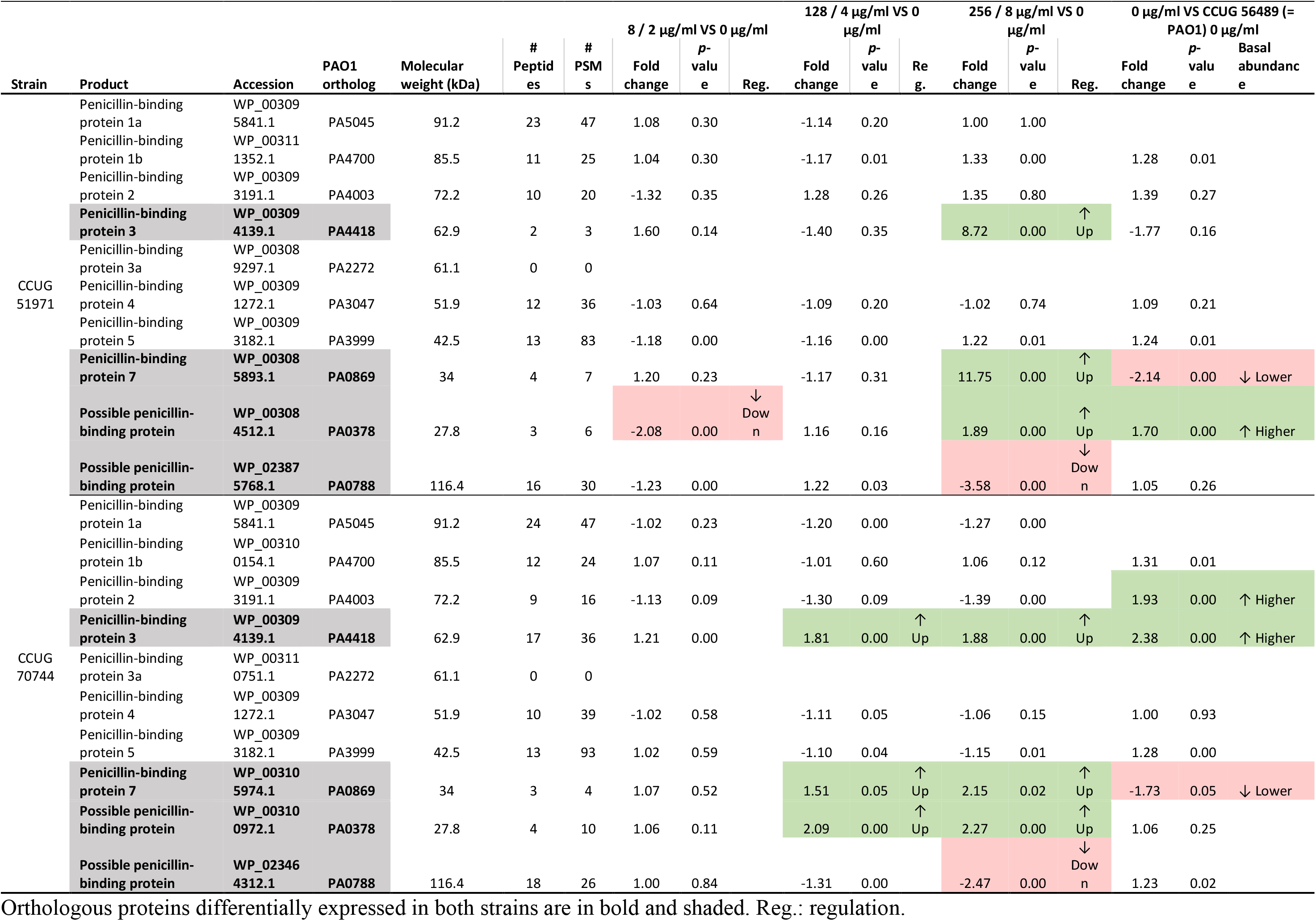
Penicillin-binding proteins (PBPs) encoded by *P. aeruginosa* CCUG 51971 and CCUG 70744 and their relative abundances.

Additionally, two possible PBPs, both containing a transglycosylase domain, were also differentially expressed. One of them (PA0378), annotated as “monofunctional biosynthetic peptidoglycan transglycosylase” by PGAP and as “probable transglycosylase” at the *Pseudomonas* Genome Database, was up-regulated similarly to PBP3 and PBP7 in both strains, i.e., up-regulated at the two highest sub-MICs in strain CCUG 70744 and only at the highest sub-MIC in strain CCUG 51971, although moderately. Strikingly, this putative PBP appeared down-regulated at the lowest sub-MIC of strain CCUG 70744. The other one (PA0788), which is annotated as PBP by PGAP, was down-regulated in both strains, at the highest sub-MIC. This might be relevant in the response to meropenem, as for instance, inactivation of PBP4 previosuly has been shown to cause β-lactam resistance by activating the CreBC two-component system and AmpC (100). Most of the other PBPs (PBP1a, PBP1b, PBP2, PBP4 and PBP5) were detected and quantitated in both strains, although none of them was differentially expressed. PBP3a was not detected in any of the strains.

### Additional differentially expressed proteins related to peptidoglycan metabolism and cell wall organization

In addition to various PBPs, several other proteins related with peptidoglycan metabolism and cell wall organization, which can modulate tolerance and resistance to β-lactams (101–103), were differentially expressed (Table 7).

**Table 7.**
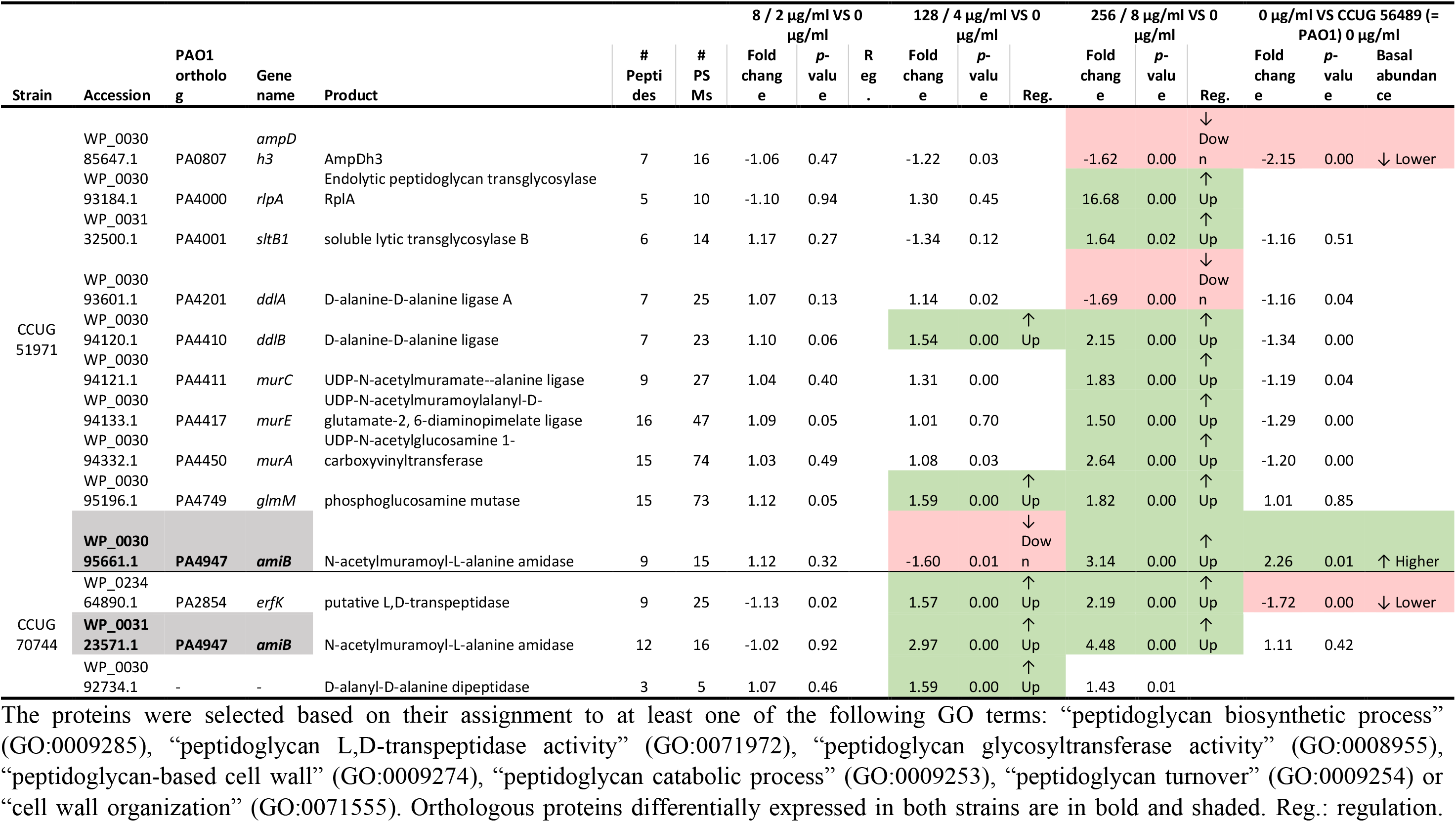
Additional differentially expressed proteins related with peptidoglycan metabolism and cell wall organization.

In strain CCUG 51971, several of these were proteins involved in the peptidoglycan biosynthesis. Among these were the D-alanine-D-alanine ligases DdlA and DdlB, which demonstrated up- and down-regulation, respectively; while DdlA appeared down-regulated only at the highest sub-MIC, DdlB was up-regulated at the two highest sub-MICs. Also, the amide ligases MurC and MurE and the enoylpyruvate transferase MurA, of the peptidoglycan biosynthesis pathway, were up-regulated at the highest sub-MIC. The phosphoglucosamine mutase GlmM, crucial for the production of the cell wall precursor UDP-N-acetylglucosamine (104), was also up-regulated, in this case, at the two highest sub-MICs.

Several proteins involved in peptidoglycan catabolism also were differentially expressed. Those included two lytic transglycosylases (RlpA and SltB1) (105, 106) which are encoded by adjacent genes in the genome and were up-regulated at the highest sub-MIC. The soluble lytic transglycosylase B (SltB1) was only slightly up-regulated, although RlpA was highly up-regulated (16.68-fold). The amidases AmpDh3 and AmiB were also differentially expressed. AmpDh3, which is one of the three AmpD homologues repressing the AmpC β-lactamase (107), was down-regulated at the highest sub-MIC, which could then be at least partially responsible for the observed up-regulation of the AmpC PDC-35. Its orthologue in strain CCUG 70744 was not detected. The other AmpD homologues (AmpD and AmpDh2) were detected but not differentially expressed, except for AmpD in strain CCUG 51971, which was not detected.

AmiB, which is crucial for cell division and its deletion has been shown to affect the outer membrane integrity, which increases its permeability and causes hyper-susceptibility to multiple antibiotics, including meropenem (108), appeared slightly down-regulated at 128 µg/ml but with clear up-regulation at 256 µg/µl. Interestingly, AmiB was also up-regulated at relatively high levels in strain CCUG 70744, in this case, at the two highest sub-MICs.

In strain CCUG 70744, only two other proteins related to peptidoglycan metabolism and cell wall organization, apart from AmiB, were differentially expressed. One was ErfK, a putative L,D-transpeptidase, possibly involved in peptidoglycan cross-linking, which was up-regulated at the two highest sub-MICs. Because of its possible L,D-transpeptidase activity, the up-regulation of ErfK could potentially compensate the harmful effects of meropenem, which blocks the D,D-transpeptidase activity of PBPs, essential for peptidoglycan cross-linking (109). The other protein was a D-alanyl-D-alanine dipeptidase, possibly involved in peptidoglycan catabolism, which was slightly up-regulated at 4 µg/µl but did not reach the significance thresholds at 8 µg/ml.

### Differentially expressed regulatory proteins

Using the webserver P2RP, a total of 599 and 597 regulatory proteins (including two-component systems, transcription factors and other DNA binding proteins) were predicted in the carbapenem-resistant strains CCUG 51971 and CCUG 70744, respectively. These represent 8.6% and 9.3% of their respective theoretical proteomes, which is in accordance with what was previously reported for strain PAO1 (28). Of all predicted regulatory proteins, 359 and 388 were detected and quantitated and, of those, 189 and 79 had significant changes in their relative abundance in at least one condition, respectively (Table 8). All regulatory proteins that were differentially expressed in at least one condition are presented in Supplementary Tables 4 and 5.

**Table 8.**
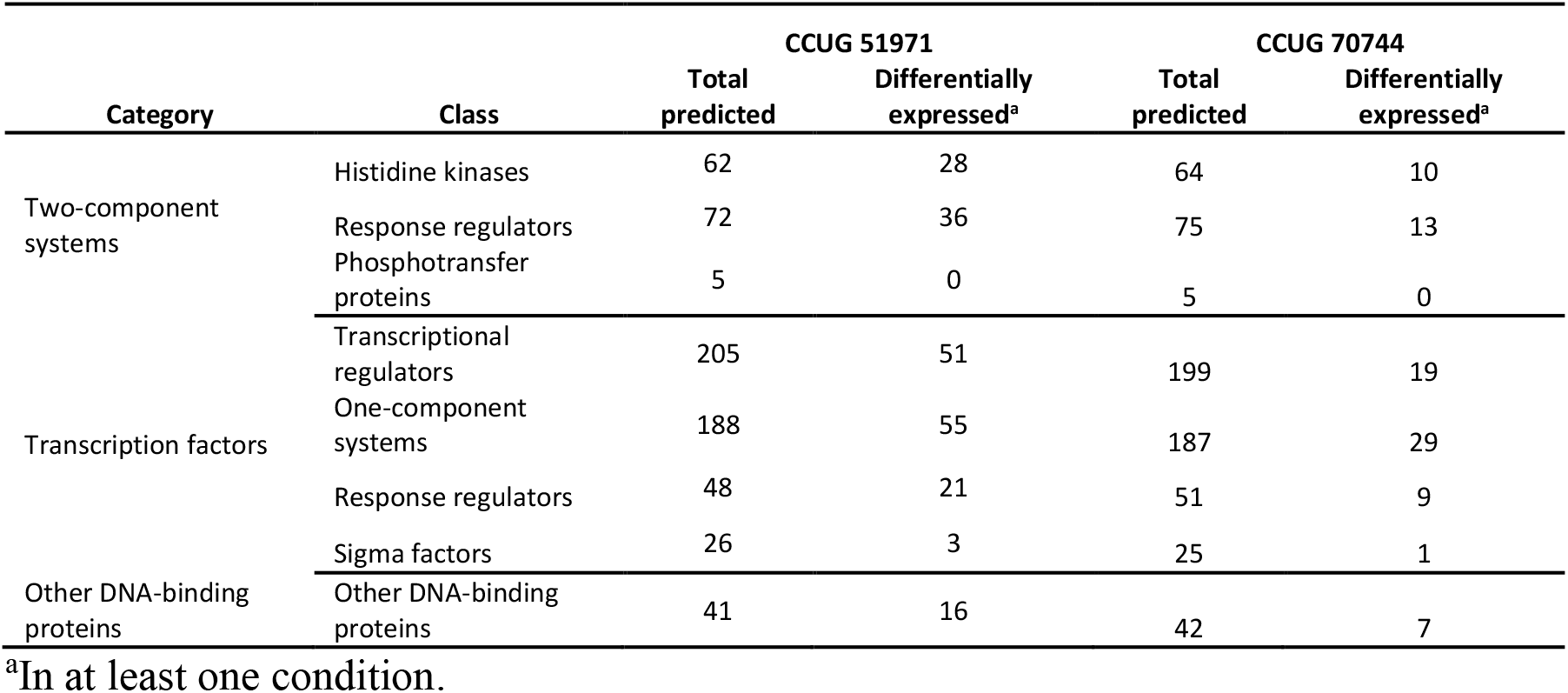
Number of regulatory proteins predicted by P2RP for each strain and number of those that were differentially expressed in at least one condition.

In both carbapenem-resistant strains, a significant proportion of regulatory proteins were differentially expressed. For instance, in strain CCUG 51971, a 45 – 50% of all the predicted two-component systems, which play a critical role in controlling how the cells respond to environmental stimuli, were differentially expressed (the majority down-regulated), most of them at 256 µg/ml. This proportion was much lower in strain CCUG 70744, but still, the large number of regulatory proteins that presented changes in their relative abundances, especially at high sub-MICs, already suggest that the responses of the cells to these sub-MICs are vast, as it has been demonstrated by the large number of differentially expressed proteins determined in both strains. This warrants future studies aiming to confirm and elucidate the role of these two-component systems in the responses of *P. aeruginosa* to carbapenems.

Several sigma factors were differentially expressed in response to meropenem. In strain CCUG 51971, AlgU, which is involved in alginate biosynthesis and thus conversion to mucoid phenotype (110), and in resistance to oxidative and heat-shock stress (111), was up-regulated at 256 µg/ml (1.63-fold). The sigma factor SbrI, which controls swarming motility and biofilm formation (112), appeared moderately down-regulated at 128 ng/µl (-1.63-fold), but below although close to the threshold of significance at 256 µg/ml (- 1.48-fold). A third differentially expressed sigma factor was FliA, which controls the synthesis of flagellin (the main component of flagella) (113) and was down-regulated at 256 µg/ml (-2.52-fold). In strain CCUG 70744, FiuI, which is involved in ferrichrome-mediated iron up-take (114), was up-regulated at 4 µg/ml (1.55-fold) but not at 8 µg/ml.

Among all predicted regulatory proteins, we specifically looked at AmpR (PA4109), a transcription factor that, in addition to proteases, quorum sensing and virulence factors, regulates the naturally-ocurring AmpC β-lactamase (115), which was up-regulated in both strains. In strain CCUG 70744, AmpR was detected and quantitated, but not differentially expressed in any condition, even though AmpC was highly up-regulated at 128 and 256 µg/ml. Comparison with strain PAO1 showed no differences in the basal abundances of AmpR in strain CCUG 70744. In strain CCUG 51971, AmpR was detected but not quantitated, which might be due to low protein abundance.

Also worth highlighting is the fact that CreD, which is the effector protein of the CreBC two-component system and plays a major role in bacterial fitness and biofilm formation, especially in the presence of β-lactam antibiotics (116), was up-regulated in all three conditions in strain CCUG 70744. In strain CCUG 51971, CreD was up-regulated at high levels (6.03-fold) but only at the highest sub-MIC. The activation of CreBC, reflected by CreD, has been previously shown to be triggered by the inhibition of PBP4 (100), which is one of the PBP for which meropenem has its highest affinity (96), and to be a major β-lactam resistance driver (100). It is also worth noting that, although the activation of a two-component system such as CreBC does not necessarily imply changes in its relative abundance, CreB was slightly down-regulated at the highest concentration in strain CCUG 70744 (-1.55-fold) and CreC was below but close to the threshold of significance (-1.47-fold), which may also have implications in the response of the bacterium to high sub-MICs of meropenem.

### Differentially expressed proteins of unknown function

Despite *P. aeruginosa* being a well characterized bacterium, the carbapenem-resistant strains CCUG 51971 and CCUG 70744 encode 1,139 and 1,251 hypothetical proteins (according to the PGAP v4.9 and v4.5 annotations available in RefSeq), respectively (i.e., 17.6 and 19.6% of the theoretical proteomes). Hundreds of these hypothetical proteins (307 and 345, respectively) were detected by the nano-LC-MS/MS analyses, which implies that they are not hypothetical, and that, from now on, should be considered “proteins of unknown function”. Additionally, both strains also encode numerous proteins with domains of unknown function (DUF) (341 and 362), of which 143 and 142 were detected, respectively.

Interestingly, many of these proteins of unknown function were also up- or down-regulated in at least one condition (244 in strain CCUG 51971 and 152 in strain CCUG 70744), suggesting that they are relevant in the response of these strains to meropenem and that they deserve further investigation to elucidate their functions and possible relation with antibiotic resistance. Moreover, 41 and 46 of them were up- or down-regulated in at least two sub-MICs, respectively. All proteins annotated as “hypothetical proteins” or containing DUFs that were differentially expressed in at least one condition are presented together with their functional annotation (assignment to COG categories, GO terms and Pfam families) in Supplementary Tables 6 and 7.

### Up-regulation of a type VI secretion system

The genome of *P. aeruginosa* harbors three T6SSs, named H1-T6SS to H3-T6SS (117), which are bacterial multiprotein devices capable of targeting effectors into prokaryotic and eukaryotic cells (118). Remarkably, all of the 14 described core components and several other associated proteins of the H1-T6SS, were up-regulated in the carbapenem-resistant strain CCUG 51971 at the highest sub-MIC (256 µg/ml) (Figure 3). These also included five effector proteins predicted for this system by the software SecReT6: the spike-carriers VgrG1 and VgrG1b and the toxins Tse3, Tse5 and Tse6. Interestingly, four of these effectors were also up-regulated at 128 and two also at 8 ng/µl. Although these toxins target bacterial cells (117, 119, 120), the H1-T6SS also has been shown to be active and important for the fitness of *P. aeruginosa* during chronic infections (121, 122), which implies that the up-regulation of this system in response to certain sub-MICs of meropenem may have direct implications in the virulence of the strain, either by outcompeting other cells or by also directly affecting also the host, as has been shown for other T6SSs of *P. aeruginosa* (123).

**Figure 3.**
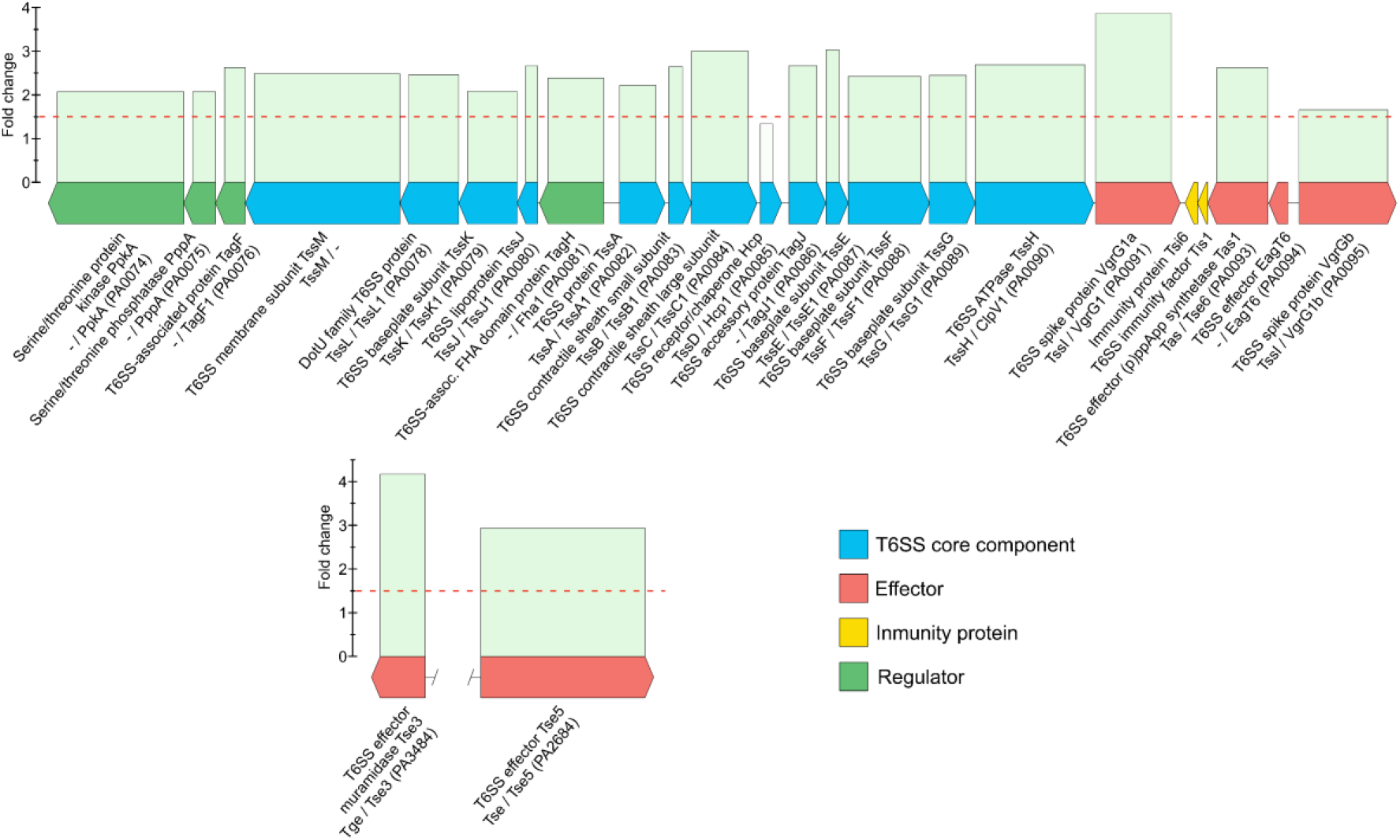
Genetic map of the H1-T6SS of *P. aeruginosa* CCUG 51971 and relative abundances of its protein products at 256 µg/ml (½ MIC) compared to 0 µg/ml. Annotations from RefSeq, SecReT6 and *Pseudomonas* Genome Database are indicated, if available.

Additionally, the H1-T6SS has been associated with decreased susceptibility to antibiotics in biofilms (124) and most of the proteins of the H1-T6SS of strain CCUG 51971 also had a higher basal relative abundance than their homologues in PAO1, suggesting that this system could potentially play a role in the resistance of this strain to multiple antibiotics.

In strain CCUG 70744, 11 of the core components of the H1-T6SS were detected and eight were quantitated, but only one was significantly up-regulated (TssB, PA0083) together with an effector protein (EagT6, PA0094). This suggests that in this strain the system is expressed but that it might only be subtly or not up-regulated in response to sub-MICs of meropenem.

### Clusters of Orthologous Groups (COG) enrichment analysis

Proteins that were differentially expressed in the cultures with the two highest sub-MICs of meropenem (i.e., ½ and ¼ of the MIC), were classified into COG categories, to determine if any category was significantly up- or down-regulated and/or significantly enriched.

The analysis revealed that, in both carbapenem-resistant strains, most of the differentially expressed proteins within the category “Translation, ribosomal structure and biogenesis” were up-regulated (Figure 4). Up-regulation of genes associated with translation and ribosomal proteins has been previously reported in response to antimicrobial agents, suggesting an increase in protein production as a protective response (125, 126). In strain CCUG 51971, most proteins within the category “Transcription” were down-regulated, although several RNA polymerase subunits were up-regulated (RpoB, RpoC and RpoZ).

**Figure 4.**
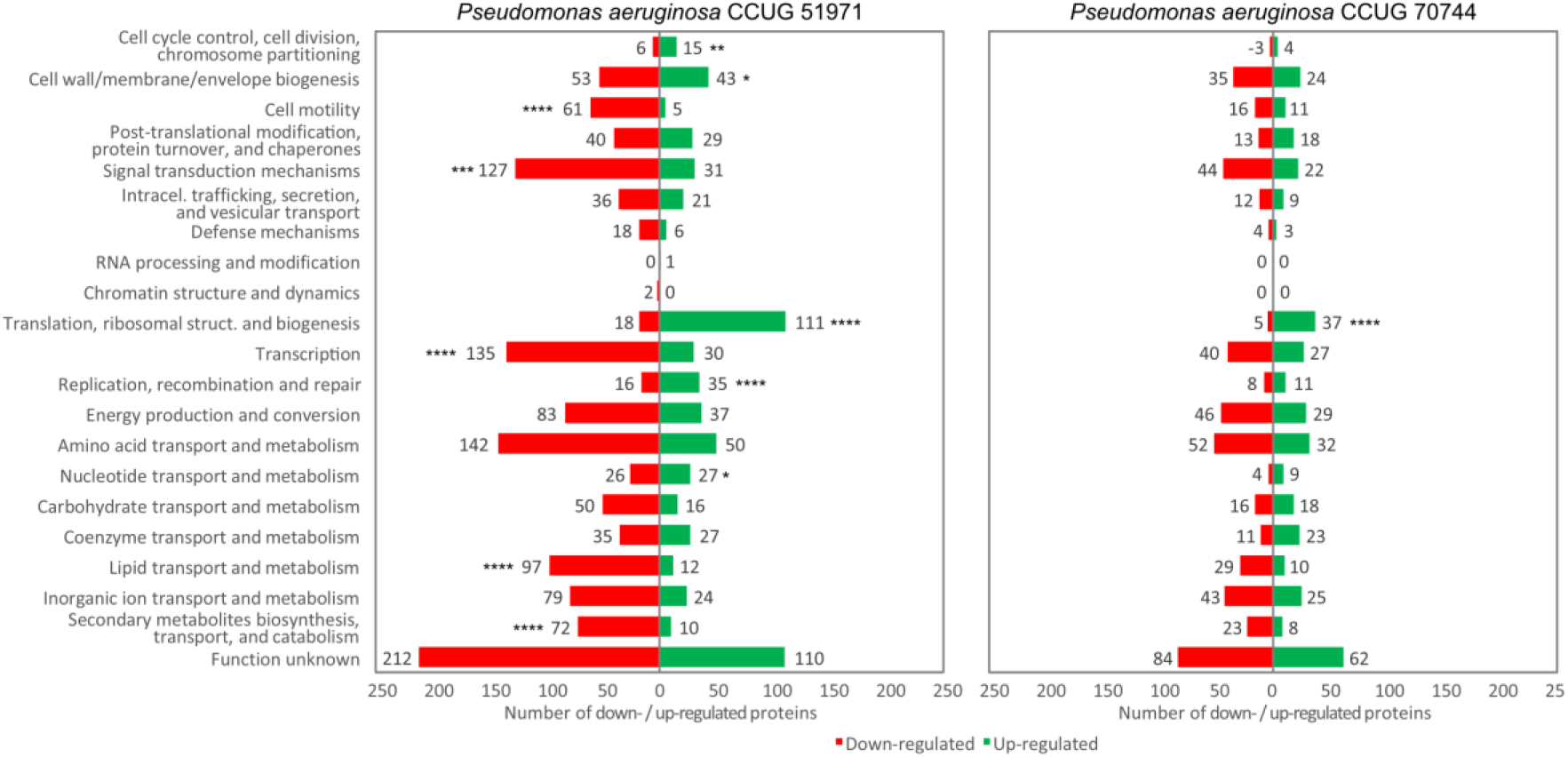
Functional classification (into COG categories) of the proteins that were up- or down-regulated in the samples cultivated with the highest sub-MIC of *P. aeruginosa* CCUG 51971 (256 µg/ml) and CCUG 70744 (8 µg/ml). Significantly up- or down-regulated COG categories are marked as follows: * if *p*-value < 0.05; ** if *p*-value < 0.005; *** if *p*-value < 0.001; **** if *p*-value < 0.0001.

In strain CCUG 70744, no other categories were significantly up- or down-regulated. However, in strain CCUG 51971, most of the differentially expressed proteins in the following COG categories were up-regulated: “Nucleotide transport and metabolism”, “Replication, recombination and repair” (which was also up-regulated at 128 µg/ml), “Cell cycle control, cell division, chromosome partitioning” and “Cell wall/membrane/envelope biogenesis”. Meanwhile, most of the differentially expressed proteins within the following categories were down-regulated: “Signal transduction mechanisms”, “Lipid transport and metabolism”, “Secondary metabolites biosynthesis, transport, and catabolism” and “Cell motility”. Of these, the last three were also significantly down-regulated at 128 µg/ml.

Additionally, the analysis in strain CCUG 51971 revealed that, when compared with all quantitated proteins, the COG categories “Signal transduction mechanisms”, “Translation, ribosomal structure and biogenesis”, “Secondary metabolites biosynthesis, transport, and catabolism” and “Cell motility” were significantly enriched (i.e., had a higher proportion of differentially expressed proteins) at 256 µg/ml. Additionally, “Cell motility” was also enriched at 128 µg/ml. In strain CCUG 70744, none of those categories was enriched, but “Energy production and conversion” and “Inorganic ion transport and metabolism” were significantly enriched at 8 µg/ml.

### Gene Ontology (GO) terms and pathway enrichment analyses

To determine if more specific subsets of proteins were significantly enriched, proteins were assigned to GO terms, KEGG pathways were annotated and GSEAs were performed. In the cases of both carbapenem-resistant strains, numerous GO terms were significantly up- or down-regulated at the highest sub-MIC and multiple up-regulated GO terms were related with translation and with ribosomal assembly (Figure 5), including those with the highest normalized enrichment scores (NES), supporting the results of the COG enrichment analysis. Indeed, Ribosome (KO03010) was the up-regulated pathway with the highest NES in both strains. Additionally, several cell wall metabolism and outer membrane-related GO terms and pathways were significantly enriched in both strains, which could be expected considering that the cell wall is the main target of β-lactam antibiotics. Moreover, the lipopolysaccharide core region biosynthetic process (GO:0009244) was up-regulated in both strains at the highest sub-MIC, which may have important implications in the virulence of the cells during infection (127). Moreover, several transport-related GO terms were down-regulated in both strains, but more abundantly in strain CCUG 70744, which reflects a response to decrease the permeability of the cells, a well-known protection mechanism against carbapenems (19). In both strains, signaling receptor activity appeared down-regulated, although most of these proteins were associated with chemotaxis and reception of siderophores. Indeed, siderophore uptake transmembrane transporter activity (GO:0015344) and iron coordination entity transport (GO:1901678) were also down-regulated with most core enrichment sequences being siderophore receptors, which could potentially be a protective response to prevent the intake of sideromycins. This may have important metabolic implications, as iron is critical for bacterial growth and survival but, certain conditions such as oxidative stress can lead to intracellular iron release and this to generation of cytotoxic reactive oxygen species (128). Additionally, chemotaxis appeared down-regulated in strain CCUG 51971, at the two highest sub-MICs. Indeed, 29 of the 44 proteins assigned to Chemotaxis (GO:0006935) were down-regulated at the highest sub-MIC, a response which probably aims to reduce mobility and promote biofilm formation. This response is in line with the down-regulation of the sigma factors SbrI and FliA, which are associated with motility (112, 113). Moreover, in strain CCUG 51971, various stress-related GO terms (response to temperature stimulus, radiation, DNA recombination, DNA repair) were up-regulated at the highest sub-MIC, suggesting that the highest sub-MIC triggered a strong stress response in this strain. Indeed, this is reflected by the up-regulation of several prophage-associated proteins, which might indicate that prophage induction was triggered by 256 µg/ml. It is also in line with the up-regulation of the sigma factor AlgU, which is associated with alginate production and resistance to oxidative and heat-shock stress (110, 111). A previous study showed that exposure of *P. aeruginosa* biofilms to imipenem induced alginate biosynthetic genes, which may be an adverse consequence of imipenem treatments (70). However, proteins of those genes were not detected in this study.

**Figure 5.**
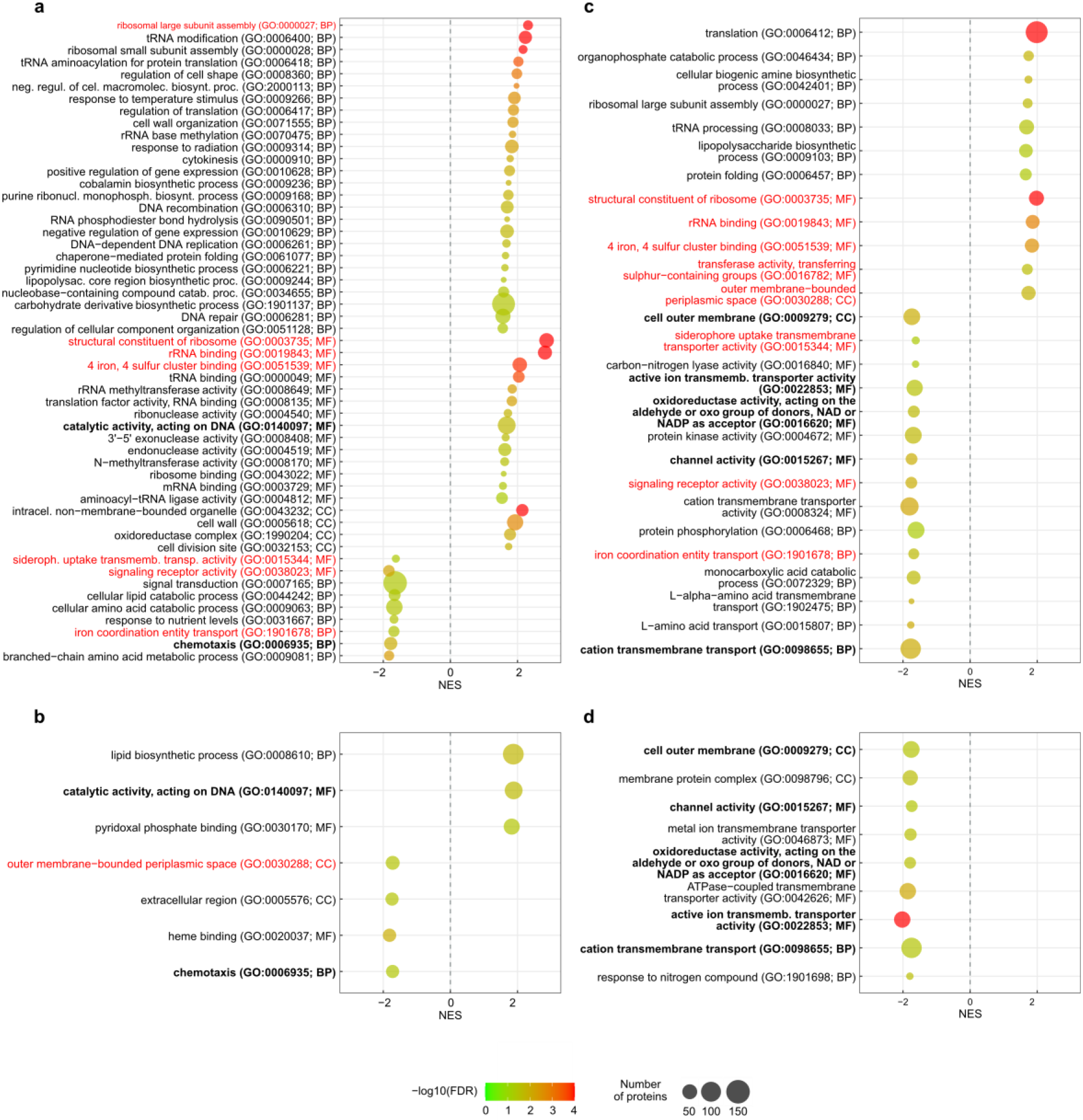
GO terms that were significantly enriched when the strains were cultivated with the two highest sub-MICs of meropenem. *P. aeruginosa* CCUG 51971 was cultivated with 256 µg/ml (**a**) and 128 µg/ml (**b**). *P. aeruginosa* CCUG 70744 was cultivated with 8 µg/ml (**c**) and 4 µg/ml (**d**). In red: GO terms enriched in both strains. In bold: GO terms enriched in both conditions of one strain. BP: biological process; MF: molecular function; CC: cellular component; FDR: false discovery rate; NES: normalized enrichment score.

**Figure 6.**
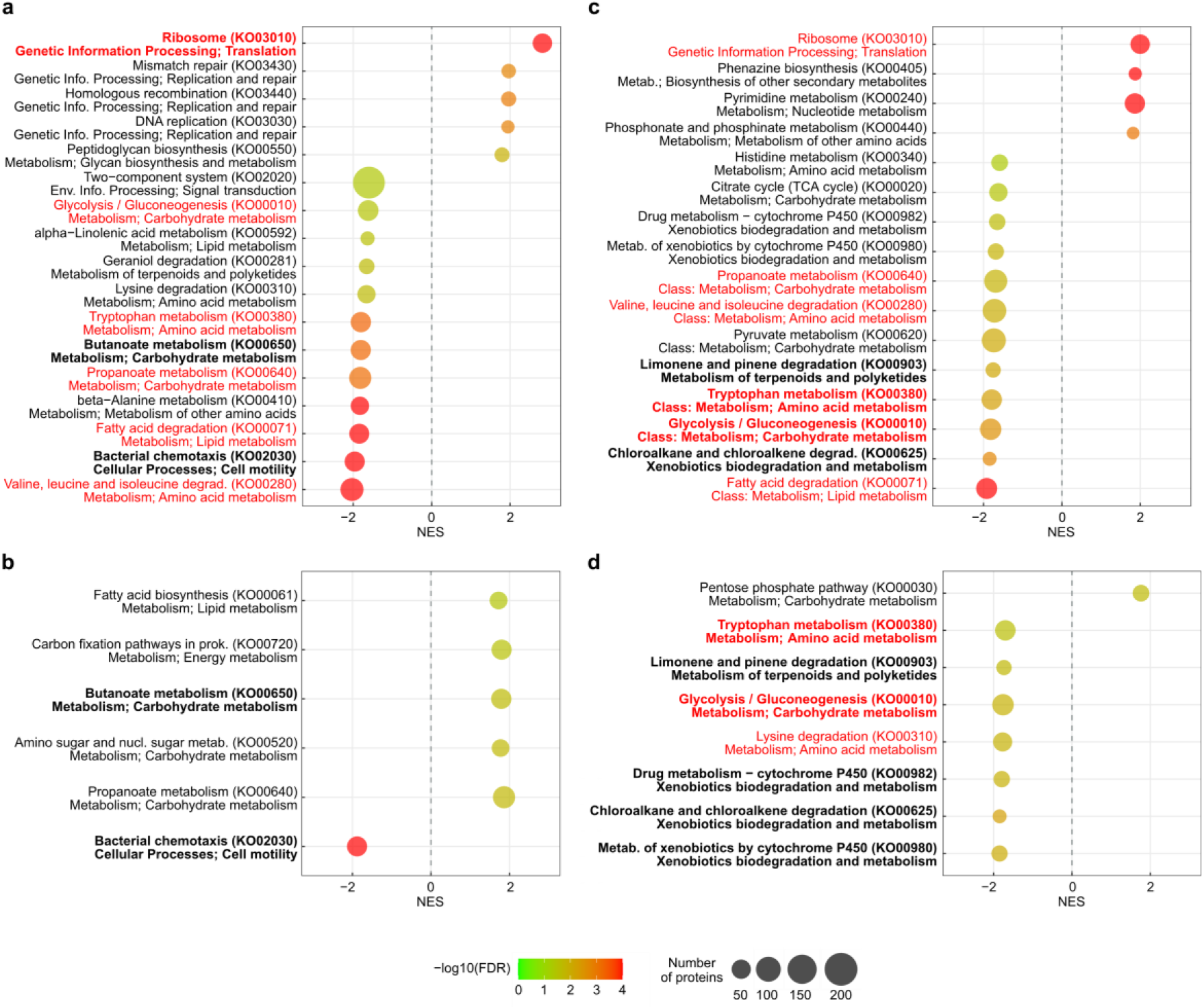
KEGG pathways that were significantly enriched when the strains were cultivated with the two highest sub-MICs of meropenem. *P. aeruginosa* CCUG 51971 was cultivated with 256 µg/ml (**a**) and 128 µg/ml (**b**). *P. aeruginosa* CCUG 70744 was cultivated with 8 µg/ml (**c**) and 4 µg/ml (**d**). The row below each pathway indicates its class. In red: pathways enriched in both strains. In bold: pathways enriched in both conditions of one strain. FDR: false discovery rate; NES: normalized enrichment score.

Interestingly, numerous metabolic pathways were significantly enriched in both strains, with most of these being down-regulated at the highest sub-MIC, affecting in both cases metabolism of carbohydrates, amino acids and lipids. Metabolic adaptation was recently shown to be able to cause antibiotic resistance (24) and in fact, multiple metabolic genes of *P. aeruginosa* have been associated with alterations in antibiotic susceptibility (129). This warrants future studies aiming to elucidate the impact of metabolic responses of *P. aeruginosa* in antibiotic resistance.

### Strengths and limitations of the study

MS-based proteomic approaches, in combination with whole-genome sequencing, offer the advantage of directly detecting produced proteins or peptides. In this study, we have shown that quantitative shotgun proteomics, using TMT labeling and nano-LC-MS/MS analysis, is an effective approach for detecting the global and specific responses of *P. aeruginosa* strains to sub-MICs of meropenem, by determining differentially expressed proteins, COG categories, GO terms and pathways. Moreover, it can be used to determine differences in the basal relative abundances of highly homologous proteins in different strains.

An inherent limitation of shotgun MS-based proteomic approaches is that, even though the techniques have advanced significantly over the last years, they still cover only a fraction of the total proteome (130), which is due to the lack of or low expression of certain genes at certain growth conditions, as well as also due to limitations of the enzymatic protein digestion (trypsin has to be capable of digesting a protein in order to generate peptides) (131), separation capacity of the liquid chromatography and inherent limitations of the mass spectrometry analysis techniques employed, such as the use of quantitative multiplexed data (132) or use of data dependent acquisition (133). Altogether these limitations can affect the coverage of an actual proteome and implies that proteins being produced and playing a significant role in the response of the strains to meropenem might have been missed due to lack of detection or quantitation. But yet, in this study, a 55.5 and 54.9% of the accessioned proteins were detected and 50.4 and 51.0% quantitated.

Transcriptomic approaches can provide expression levels for most genes being expressed and, hence, may sometimes be perceived as a more comprehensive and less cumbersome alternative. However, low correlation between transcriptomic and proteomic data has been observed and can occur due to multiple reasons (134), whereas proteins are the ultimate effectors of the organism. Thus, quantitative proteomics represents a more direct approach for observing the responses of cells, although a good approach could also be to apply multiple approaches such as shotgun genomic, transcriptomic and proteomic analyses, as proposed by Hua *et al.* (135). In any case, although many proteins might not be generated at all or produced at very low levels, we were able to detect and quantitate more than half of the theoretical proteome, which provides a comprehensive image of the response of the bacteria to sub-MICs of meropenem.

Another limitation of isobaric labeling-based approaches is that they can only detect changes in relative abundances but cannot do absolute quantification. Thus, proteins encoded by genes that have been constitutively under or overexpressed but that do not vary between conditions, will not be noted, even though those high or low levels of abundance might be involved in resistance. However, this issue can be addressed by including strains with highly similar orthologous sequences, as in this case PAO1. For instance, no changes in the relative abundance of PIB-1 nor MexXY were detected in strain CCUG 70744, but the comparison with strain PAO1 revealed that they had significantly higher basal abundances, which has previously been associated with β-lactam resistance (78, 92).

In any case, our study illustrates the potential of quantitative proteomic approaches to reveal global and specific changes in the proteomes of carbapenem-resistant *P. aeruginosa* strains with different mechanisms of resistance, in response to antibiotic stress and has revealed numerous proteins that, at least in the context of these two carbapenem-resistant *P. aeruginosa* strains, are differentially expressed when cultivated with sub-MICs of meropenem. This will serve as a guideline for future studies aiming to confirm the role of these proteins in the response of *P. aeruginosa* to meropenem. It will also serve as a basis for future investigations searching alternative resistance targets and low-level resistance mechanisms. However, an obvious limitation of this study is that only two carbapenem-resistant strains of only two high-risk clones have been used, while *P. aeruginosa* is a diverse species with an exceptionally large and open pan-genome (136). This means that the analysis of additional strains with different genetic backgrounds will probably reveal new and different responses, as it has been observed between strains CCUG 51971 and CCUG 70744.

Thus, it will be important to look at how frequent and conserved these differentially expressed proteins are among other strains of *P*. *aeruginosa*, to determine if they are relevant to only a few strains or to a larger proportion of the species. This could be done by screening the >7,000 publicly available genome sequences of *P. aeruginosa*, although further experiments confirming the involvement of these proteins in resistance to meropenem in numerous strains should be performed. Thus, future studies including more strains with different and diverse genomic backgrounds and acquired resistance mechanisms, as well as additional antimicrobial compounds and conditions (e.g., *in vivo* models), will be essential to determine how conserved are in *P. aeruginosa* the putative mechanisms revealed in this study and to elucidate further responses.

## Conclusion

The exposure of the carbapenem-resistant *P. aeruginosa* strains CCUG 51971 (ST235) and CCUG 70744 (ST395) to sub-MICs of meropenem caused significant strain- and concentration-specific changes in their proteomes. The marked responses involve complex interactions between multiple canonical and non-canonical mechanisms, each of which might be accountable at different levels for the low susceptibility of these strains to carbapenem antibiotics. Additionally, multiple proteins of unknown function were differentially expressed in both strains, in response to exposure to sub-MIC levels of meropenem, which warrants future studies to explore their function and determine if and how they are involved in the responses of these strains to meropenem. Our study demonstrates that quantitative shotgun proteomics is effective for identifying different and complex mechanisms that collectively may play a role in reduced susceptibility of *P. aeruginosa* to meropenem. Our study provides a framework for elucidating responses in expression and determining alternative mechanisms of resistance on exposure to antibiotics, that may lead to identifying new drug targets in pathogens.

## Data availability statement

The strains used in this study are available at the Culture Collection University of Gothenburg (CCUG, Gothenburg, Sweden; www.ccug.se). The complete genome sequence of *P. aeruginosa* CCUG 51971 (= PA 66) has been deposited in DDBJ/ENA/GenBank under the accession no. CP043328. The complete genome sequence of *P. aeruginosa* CCUG 70744 was already publicly available in DDBJ/ENA/GenBank under the accession no. CP023255. The Illumina and Oxford Nanopore sequence reads of *P. aeruginosa* CCUG 51971 have been deposited in the Sequence Read Archive (SRA) (137), under the accession numbers SRX6772487, SRX6772488 and SRX6772489. The mass spectrometry proteomics data has been deposited in the ProteomeXchange Consortium (138) via the PRIDE partner repository (139) with the dataset identifier PXD034987.

## Author contributions

Conceptualization – F.S-S., I.A., E.R.B.M., R.K.; Methodology – F.S-S., D.J-L., R.K.; Validation – F.S-S., D.J-L., N.P.M., I.A., and R.K.; Original draft preparation – F.S-S.; Review & Editing – F.S-S., D.J-L., N.P.M., I.A., E.R.B.M., R.K.; Supervision – E.R.B.M. and R.K.; Project administration – E.R.B.M. and R.K.; Funding acquisition – D.J-L., I.A., E.R.B.M., R.K.

## Acknowledgements

The authors thank the staff at the Culture Collection University of Gothenburg (CCUG) for maintaining and providing the strains used in this study. Anders Malmborg and the Substrate Department of the Department of Clinical Microbiology, Sahlgrenska University Hospital, are acknowledged for media preparation. The authors also acknowledge Christian G. Giske at the Karolinska Institutet for providing strain PA 66. The authors acknowledge the National Reference Laboratory for Antibiotic Resistance (Växjö, Sweden), for the determination of minimal inhibitory concentrations. Annika Thorsell and Maria Segeda, at the Proteomics Core Facility (Sahlgrenska Academy, University of Gothenburg), are acknowledged for generating the quantitative proteomic data and performing the Proteome Discoverer analyses. We also acknowledge Jari Martikainen and Björn Andersson, at the Bioinformatics Core Facility (Sahlgrenska Academy, University of Gothenburg), for constructing the heatmaps, PCA plots and COG enrichment analyses. The computations were partially performed on resources provided by the Swedish National Infrastructure for Computing (SNIC) through the Uppsala Multidisciplinary Center for Advanced Computational Science (UPPMAX) under project SNIC 2019/8-176. The authors acknowledge Anna Johnning and Erik Kristiansson at Chalmers University of Technology and Johan Bengtsson-Palme at the University of Gothenburg for valuable discussions. They also acknowledge Leonarda Achá Alarcón at the University of Gothenburg for proof-reading the manuscript.

## Abbreviations

ABC: ATP-binding cassette
ANIb: average nucleotide identity based on BLAST
bRP-LC: basic reversed-phase liquid chromatography
CARD: Comprehensive Antibiotic Resistance Database
CCUG: Culture Collection University of Gothenburg
CFU: colony forming unit.
CID: collision-induced dissociation
COG: Cluster of Orthologous Groups
DUF: domain of unknown function
EC: Enzyme Commission
EDTA: ethylenediaminetetraacetic acid
ESAC: extended-spectrum AmpC
EUCAST: European Committee on Antimicrobial Susceptibility Testing
FASP: filter-aided sample preparation
FDR: false discovery rate GO Gene Onthology
GSEA: Gene Set Enrichment Analysis
HCD: higher-energy collisional dissociation
ICE: integrative and conjugative element
KEGG: Kyoto Encyclopedia of Genes and Genomes
LC-MS/MS: liquid chromatography tandem-mass spectrometry
MBL: metallo-β-lactamase
MIC: minimal inhibitory concentration
MLST: multilocus sequence typing
NES: normalized enrichment score
PBP: penicillin-binding protein
PBS: phosphate-buffered saline
PC: principal component
PCA: principal component analysis
PDC: *Pseudomonas*-derived cephalosporinase
PGAP: Prokaryotic Genome Annotation Pipeline
PSM: peptide-spectrum match
RGI: Resistance Gene Identifier
RND: resistance-nodulation-cell division
SDC: sodium deoxycholate
SDS: sodium dodecyl sulfate
ST: sequence type
Sub-MIC: subminimal inhibitory concentration
T6SS: type VI secretion system
TEAB: triethylammonium bicarbonate
TMT: tandem mass tag

## Supplementary information

**Supplementary figure 1.**
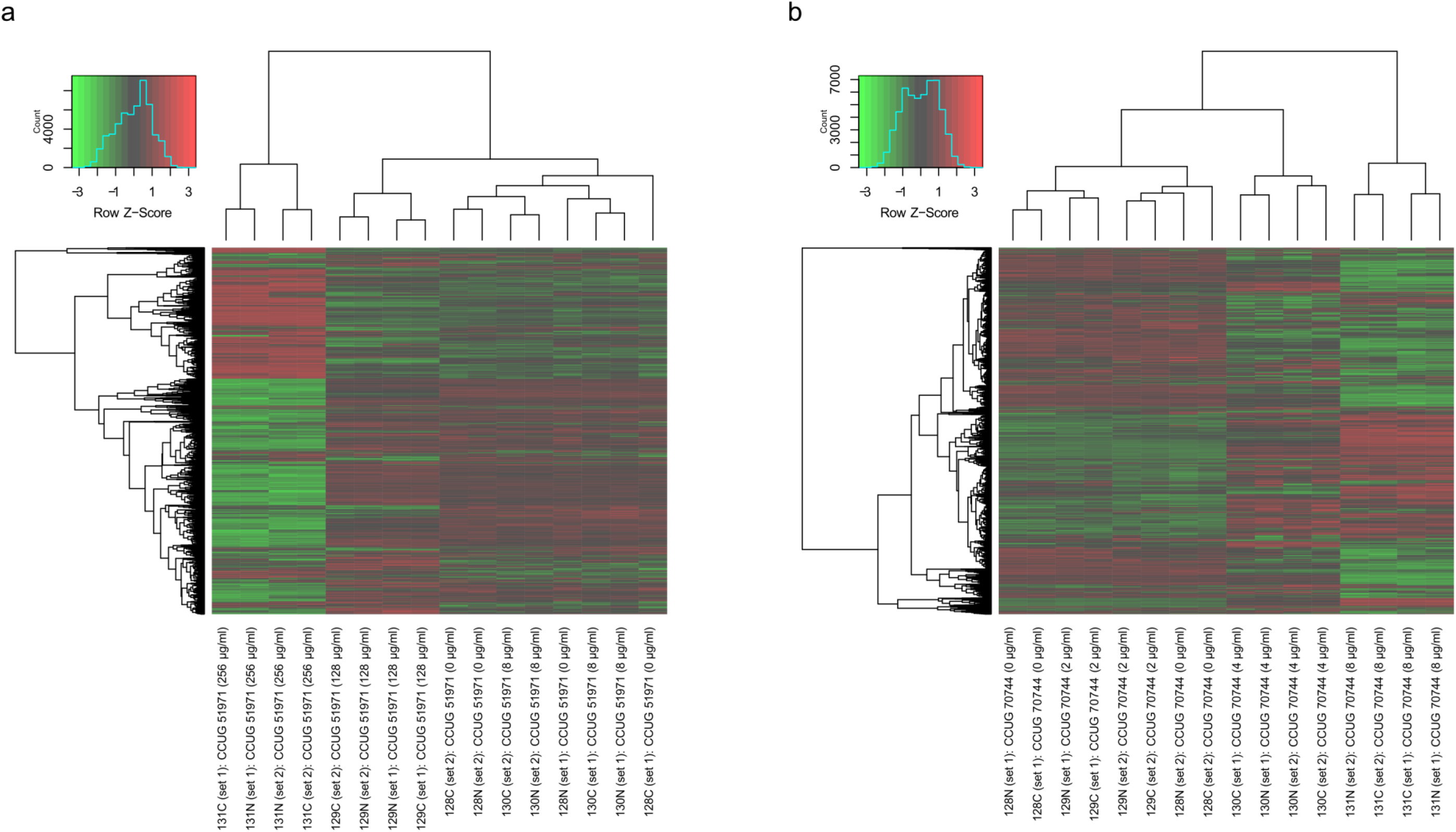
Heatmap representations and dendrograms of the quantitative nano-LC-MS/MS-based proteomic data of *P. aeruginosa* CCUG 51971 (a) and *P. aeruginosa* CCUG 70744 (b). For each strain, the four replicates of each of the four conditions are included.

**Supplementary figure 2.**
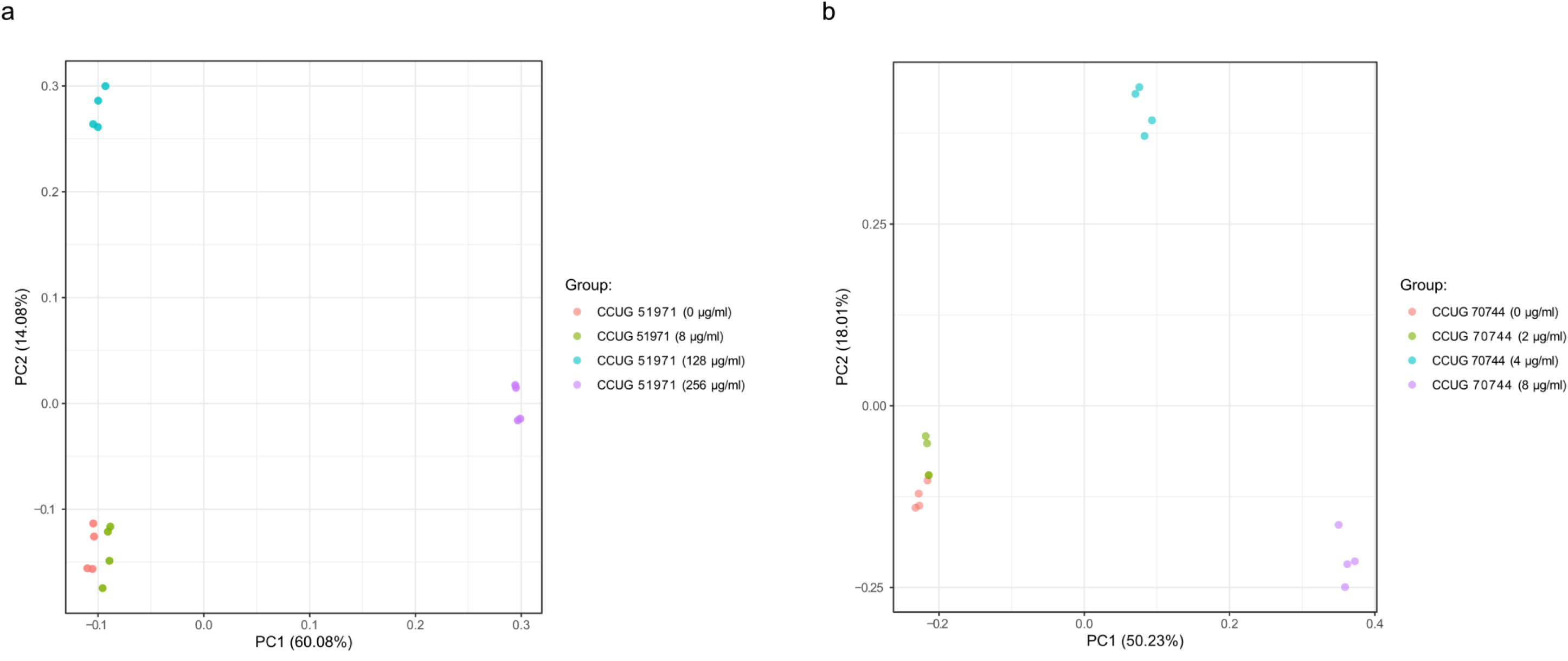
Principal component analysis (PCA) plots, based on the results of the quantitative nano-LC-MS/MS-based proteomic analyses of the quadruplicate samples of: **a** *P. aeruginosa* CCUG 51971; and **b** *P. aeruginosa* CCUG 70744, exposed to different sub-MICs of meropenem (strain CCUG 51971: 8, 128 and 256 µg/ml; strain CCUG 70744: 2, 4 and 8 µg/ml) and to no antibiotic. Each plot shows four groups of samples (one per meropenem concentration), distributed according to their similarity by principal components (PC) 1 and 2. Each dot represents one replicate.

**Supplementary figure 3.**
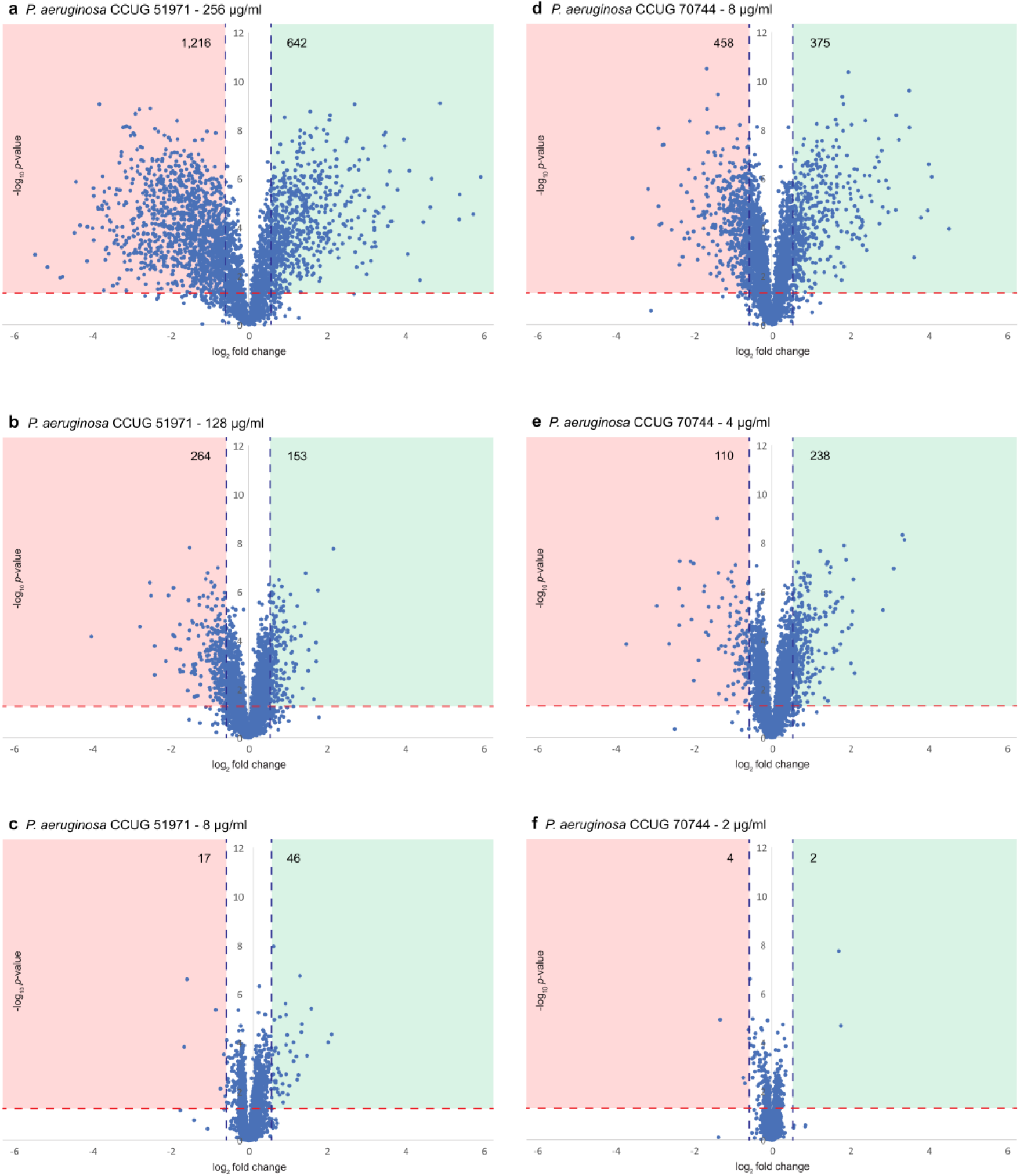
Volcano plots of the six quantitative proteomic comparisons. **a** *P. aeruginosa* CCUG 51971 at 256 µg/mL compared to 0 µg/mL. **b** *P. aeruginosa* CCUG 51971 at 128 µg/mL compared to 0 µg/mL. **c** *P. aeruginosa* CCUG 51971 at 8 µg/mL compared to 0 µg/mL. **d** *P. aeruginosa* CCUG 70744 at 8 µg/mL compared to 0 µg/mL. **e** *P. aeruginosa* CCUG 70744 at 4 µg/mL compared to 0 µg/mL. **f** *P. aeruginosa* CCUG 70744 at 2 µg/mL compared to 0 µg/mL. The x-axis shows the log_2_ of the fold changes and the y-axis shows the -log_10_ of the *p*-values. Quantitated proteins are represented by dots. The colored squares represent the areas of significance, i.e., proteins within those areas had significantly down- or up-regulated abundance levels (i.e., fold changes equal or higher than ±1.5 and *p*-values <0.05). The numbers within the colored squares indicate the number of significantly down- or up-regulated proteins.

**Supplementary table 1.** Number of proteins and peptides detected and quantitated in *P. aeruginosa* strains CCUG 51971 and CCUG 70744.

**Supplementary table 2.** List of proteins of *Pseudomonas aeruginosa* CCUG 51971 detected with more than one peptide. The table includes their relative abundances (sub-MICs vs. no antibiotic; if quantitated), basal relative abundance compared to strain PAO1 (if fulfilling the criteria for being included in the inter-strain the comparison, i.e., ≥99% of sequence identity and with 100% coverage over the longer protein), *P. aeruginosa* PAO1 orthologues and functional annotations. Orthologous proteins differentially expressed in both strains are in bold and shaded.

**Supplementary table 3.** List of proteins of *Pseudomonas aeruginosa* CCUG 70744 detected with more than one peptide. The table includes their relative abundances (sub-MICs vs. no antibiotic; if quantitated), basal relative abundance compared to strain PAO1 (if fulfilling the criteria for being included in the inter-strain the comparison, i.e., ≥99% of sequence identity and with 100% coverage over the longer protein), *P. aeruginosa* PAO1 orthologues and functional annotations. Orthologous proteins differentially expressed in both strains are in bold and shaded.

**Supplementary table 4.** List of regulatory proteins of *Pseudomonas aeruginosa* CCUG 51971 that were differentially expressed in at least one sub-MIC. The table includes their relative abundances (sub-MICs vs. no antibiotic; if quantitated), basal relative abundance compared to strain PAO1 (if fulfilling the criteria for being included in the inter-strain comparison, i.e., ≥99% of sequence identity and with 100% coverage over the longer protein), *P. aeruginosa* PAO1 orthologues and functional annotations. Orthologous proteins differentially expressed in both strains are in bold and shaded.

**Supplementary table 5.** List of regulatory proteins of *Pseudomonas aeruginosa* CCUG 70744 that were differentially expressed in at least one sub-MIC. The table includes their relative abundances (sub-MICs vs. no antibiotic; if quantitated), basal relative abundance compared to strain PAO1 (if fulfilling the criteria for being included in the inter-strain comparison, i.e., ≥99% of sequence identity and with 100% coverage over the longer protein), *P. aeruginosa* PAO1 orthologues and functional annotations. Orthologous proteins differentially expressed in both strains are in bold and shaded.

**Supplementary table 6.** List of proteins of *Pseudomonas aeruginosa* CCUG 51971 annotated as “hypothetical proteins” or as proteins with domain of unknown function in RefSeq that were differentially expressed in at least one sub-MIC. The table includes their relative abundances (sub-MICs vs. no antibiotic; if quantitated), basal relative abundance compared to strain PAO1 (if fulfilling the criteria for being included in the inter-strain comparison, i.e., ≥99% of sequence identity and with 100% coverage over the longer protein), *P. aeruginosa* PAO1 orthologues and functional annotations. Orthologous proteins differentially expressed in both strains are in bold and shaded.

**Supplementary table 7.** List of proteins of *Pseudomonas aeruginosa* CCUG 70744 annotated as “hypothetical proteins” or as proteins with domain of unknown function in RefSeq that were differentially expressed in at least one sub-MIC. The table includes their relative abundances (sub-MICs vs. no antibiotic; if quantitated), basal relative abundance compared to strain PAO1 (if fulfilling the criteria for being included in the inter-strain comparison, i.e., ≥99% of sequence identity and with 100% coverage over the longer protein), *P. aeruginosa* PAO1 orthologues and functional annotations. Orthologous proteins differentially expressed in both strains are in bold and shaded.

**Supplementary table 8.** Proteins of the H1-T6SS of *P. aeruginosa* CCUG 51971. The table includes their relative abundances (sub-MICs vs. no antibiotic; if quantitated), basal relative abundance compared to strain PAO1 (if fulfilling the criteria for being included in the inter-strain comparison, i.e., ≥99% of sequence identity and with 100% coverage over the longer protein), *P. aeruginosa* PAO1 orthologues and functional annotation.

